# Butyrolactol A potentiates caspofungin efficacy against resistant fungi via phospholipid flippase inhibition

**DOI:** 10.1101/2025.01.06.630955

**Authors:** Xuefei Chen, H. Diessel Duan, Michael J. Hoy, Kalinka Koteva, Michaela Spitzer, Allison K. Guitor, Emily Puumala, Aline A. Fiebig, Guanggan Hu, Bonnie Yiu, Sommer Chou, Zhuyun Bian, Yeseul Choi, Amelia Bing Ya Guo, Wenliang Wang, Sheng Sun, Nicole Robbins, Anna Floyd Averette, Michael A. Cook, Ray Truant, Lesley T. MacNeil, Eric D. Brown, James W. Kronstad, Brian K. Coombes, Leah E. Cowen, Joseph Heitman, Huilin Li, Gerard D. Wright

## Abstract

Fungal infections cause millions of deaths annually and are challenging to treat due to limited therapeutic options and rising resistance. Cryptococci are intrinsically resistant to the latest generation of antifungals, echinocandins, while *Candida auris*, a notorious global threat, is also increasingly resistant. We performed a natural product extract screen to rescue caspofungin fungicidal activity against *Cryptococcus neoformans* H99 and identified butyrolactol A, which restores echinocandin efficacy against resistant fungal pathogens, including multidrug-resistant *C. auris*. Mode of action studies revealed that butyrolactol A inhibits the phospholipid flippase Apt1–Cdc50, blocking phospholipid transport. Cryo-electron microscopy analysis of the Apt1•butyrolactol A complex revealed that the flippase is trapped in a dead-end state. Apt1 inhibition disrupts membrane asymmetry, vesicular trafficking, and cytoskeletal organization, thereby enhancing echinocandin uptake and potency. This study identifies lipid flippases as promising antifungal targets and demonstrates the potential of revisiting natural products to expand the antifungal arsenal and combat resistance.

## Introduction

Fungal infections affect over a billion people worldwide and are responsible for an estimated 3.8 million deaths annually, with 2.5 million directly attributable to fungal disease,^1, 2^ exceeding tuberculosis and malaria.^3, 4^ The World Health Organization and Centers for Disease Control and Prevention have raised global concern over the rise of multidrug-resistant fungal pathogens, such as *Candida auris*, which poses a serious and growing health threat.^5^ Among the most critical fungal pathogens are species of the genus *Cryptococcus*, which cause life-threatening meningitis, pneumonia, and other central nervous system diseases.^6, 7^ Cryptococcal meningitis is the most prevalent disseminated fungal disease among people with AIDS, responsible for approximately 15% of AIDS-related deaths annually (>181,000), with mortality rates approaching 100% if infections remain untreated.^8, 9^

Despite the substantial impact of fungal infections on human health, treatment options are limited to only three major drug classes approved for clinical use.^8, 10^ Polyenes such as amphotericin B (AMB), which bind the fungal membrane sterol ergosterol, remain a frontline therapy for cryptococcosis, but their use is constrained by severe toxicity, including hemolysis.^11^ Azoles are another commonly used drug class that targets ergosterol biosynthesis but generally exerts fungistatic effects, causing selective pressure for resistance development.^12, 13^ Though the pyrimidine antimetabolite 5-fluorocytosine can be effectively paired with azoles or AMB, its monotherapy is precluded by the rapid emergence of resistance and lack of availability in many low-resource settings.^14, 15^ The most recent antifungal class, echinocandins, is ineffective against *Cryptococcus* species, and resistant *C. auris* strains are increasingly prevalent.

Echinocandins, such as the semi-synthetic drug caspofungin (Figure 1A), inhibit the β- (1,3)-D-glucan synthase subunit Fks1, which polymerizes the essential polysaccharide component of most fungal cell walls, resulting in fungal cell lysis.^16, 17^ The absence of cell walls in mammalian cells has made echinocandins highly successful, low-toxicity antifungal drugs.^18^ ^19^ In *C. neoformans,* the *FKS1* gene is essential for fungal viability, and the purified cryptococcal β-(l,3)-D-glucan synthase is inhibited by echinocandins *in vitro*.^20, 21^ Nevertheless, cryptococci are intrinsically resistant to echinocandin therapy in part due to the action of cell division control protein 50 (CDC50), the noncatalytic subunit of a binary lipid flippase complex.^22, 23^ Targeting the extracellular loop of Cdc50 with an antifungal peptide modestly enhances caspofungin activity with a fractional inhibitory concentration index (FICI) of 0.5, demonstrating a potential strategy to sensitize *Cryptococcus* to caspofungin through CDC50 inhibition.^24^

**Figure 1.**
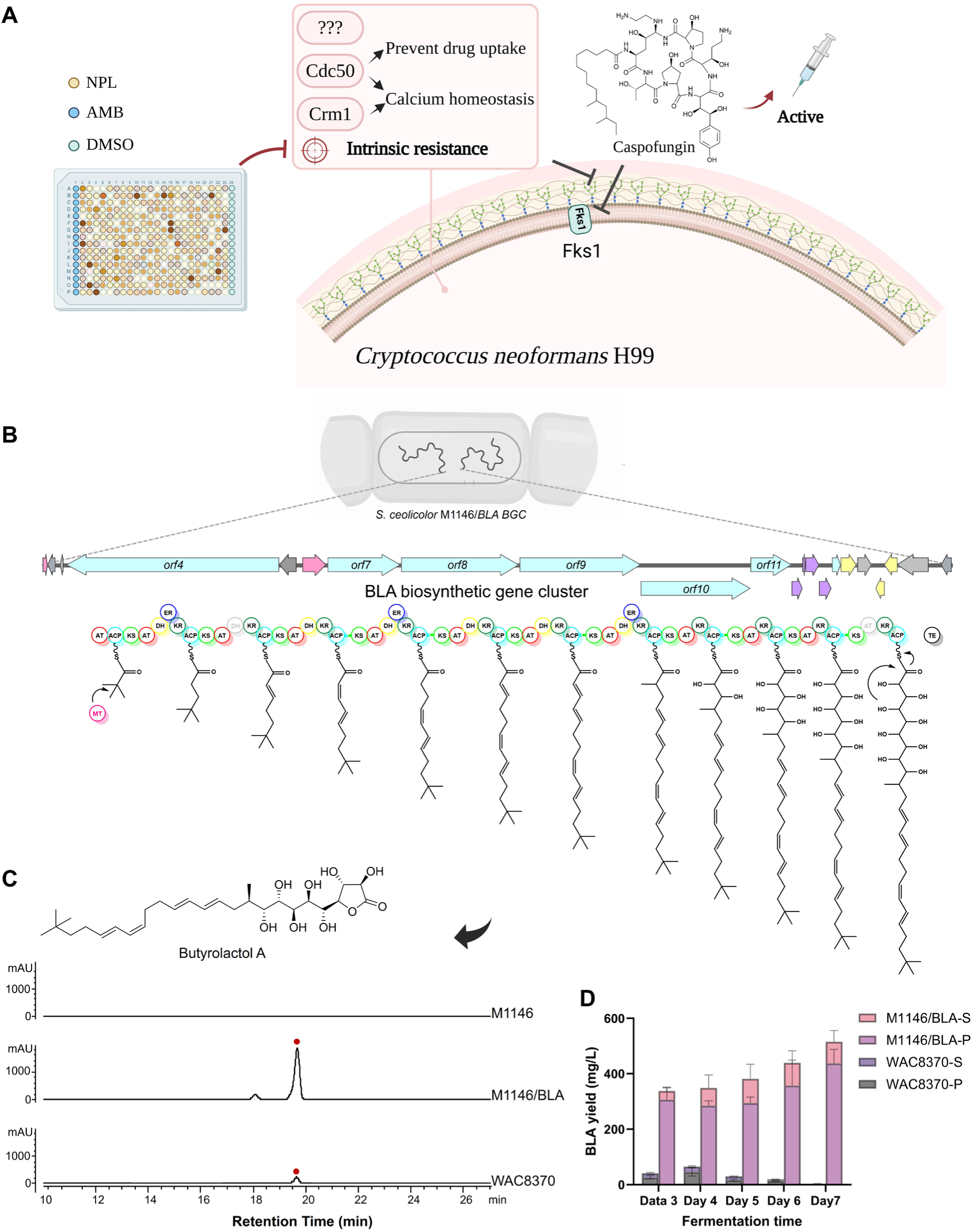
A high throughput screen for potentiators of caspofungin against *Cryptococcus neoformans* H99 and generation of high-yield biosynthetic producers of butyrolactol. **A.** (A) Overview of the screening strategy using the natural product library (NPL) to identify caspofungin potentiators that can overcome intrinsic resistance in *Cryptococcus* species. Amphotericin B (AMB) is a positive antifungal control. See also Figure S1A. **(B)** Identification and expression of the butyrolactol A (BLA) biosynthetic gene cluster (BGC) from the genome of *Streptomyces ardesiacus* WAC8370. The organization of BLA BGC is color-coded: blue for polyketide backbone (PKS), purple for hydroxymalonyl-ACP biosynthesis genes, pink for the transcriptional regulator, light yellow for the transporter, and grey for other genes. PKS domains are labeled as follows: AT, acyltransferase; KS, keto synthase; ACP, acyl carrier protein; KR, ketoreductase; DH, dehydratase; ER, enoylreductase; TE, thioesterase. See also Table S4. (C) Chromatographic analysis of BLA produced by heterologous hosts, compared to the natural producer *S. ardesiacus* WAC8370. M1146 refers to *S. coelicolor* M1146; M1146/BLA contains the butyrolactol A biosynthetic gene cluster. See also Figure S3. (D) Refactoring and heterologous expression significantly enhanced the production of butyrolactol A over time. Extracellular and intracellular levels of butyrolactol A were quantified in wild-type *S. ardesiacus* WAC8370 and the engineered *S. coelicolor* strain (M1146/BLA) from day 3 to day 7 of fermentation. S = supernatant (conditioned media); P = pellet (cell-associated fraction).

Identifying adjuvants that reverse echinocandin resistance presents an opportunity to repurpose this drug class for cryptococcal infections and to rescue its efficacy against other resistant pathogens.^25^ Reasoning that natural products offer an untapped source of echinocandin potentiators, we screened a subset of our collection of microbial natural product extracts^26^ for compounds that sensitize *C. neoformans* to caspofungin. Here, we report the discovery of butyrolactol A, a polyketide antifungal natural product that unexpectedly enhances echinocandin activity against cryptococci and multidrug-resistant *Candida* spp. by inhibiting the Apt1–Cdc50 phospholipid flippase. This previously unrecognized mechanism of action expands the antifungal landscape and enriches our limited antifungal arsenal with an orthogonal combination strategy to combat drug-resistant fungal infections.

## Results

### A high throughput screen for potentiators of caspofungin against *C. neoformans* H99 identifies butyrolactol A

Approximately 4,000 methanolic extracts derived from our in-house collection of actinomycetes and fungi^26^ were screened against *C. neoformans* H99 in the presence and absence of 4 µg/mL caspofungin (1/8 MIC) (Figures 1A, S1A). Methanol was selected as the extraction solvent due to its broad polarity range and compatibility with downstream analytical workflows, allowing for the efficient recovery of diverse microbial metabolites for screening while excluding highly hydrophobic metabolites.

Six extracts exhibited robust and reproducible caspofungin-potentiating capacity in a dose-dependent manner (Figure S1B). Among these, the anticancer compound actinomycin D, known for its cytotoxicity in mammalian cells, was identified from strains WAC237 and WAC1315 as an active caspofungin adjuvant (Figures S1C-S1D). The remaining four isolates, WAC205, WAC6266, WAC6818, and WAC8370, all yielded a shared potentiator, compound **1**, which displayed a UV absorption maximum at 236 nm in acetonitrile (Figures S1E and S1F). Structural elucidation of compound **1** by high-resolution mass spectrometry (HRMS) and multidimensional NMR spectroscopy, identified it as butyrolactol A (BLA), an antifungal polyketide polyol first reported in 1992 from *Streptomyces rochei* S785-16 (Figure S1G, Supplementary Note 1).^27^As the absolute stereochemistry of BLA remained undefined and was potentially relevant to its bioactivity and target engagement, we undertook its stereochemical assignment. The configurations at C-2 and C-3 were determined to be R, R by Mosher ester analysis using (R)- and (S)-MTPA-Cl (Figure S2, Tables S1-S3). Additional stereocenters in the γ-lactone ring and polyol tail were assigned as 3R, 4S, 5R, 6R, 7S, 8S, 9R, and 10R based on scalar coupling constants (J values), ROESY correlations, and HSQC-HECADE spectra (Supplementary Notes 1 and 2). BLA also exhibited a specific optical rotation of –30.2° (c = 0.1, 10% DMSO in methanol, 25 °C), consistent with the presence of a single enantiomer.

### A synthetic biology strategy to develop high-yield producers of butyrolactol A

To address the poor and inconsistent yield of BLA from the native producer *S. ardesiacus* WAC8370, a common challenge for secondary metabolite production under laboratory fermentation conditions potentially due to tight transcriptional regulation that hampers downstream investigations,^28^ we employed a synthetic biology strategy to establish a robust heterologous expression system. Whole-genome sequencing revealed a BLA biosynthetic gene cluster (BGC) highly similar to the previously characterized type I polyketide synthase system from *Streptomyces* sp. NBRC 110030 (Figure S3A, Table S4).^29^ Due to its size and restriction enzyme site availability, the BGC was cloned in two overlapping segments. A 71,401-bp genomic fragment encompassing the core biosynthetic and tailoring genes was captured using transformation-associated recombination (TAR) and introduced into the model chassis *Streptomyces coelicolor* M1146 for heterologous expression (Figures 1B, S3B-S3C, Table S4).^30^ To ensure complete pathway reconstitution, the downstream operon (*orf19*–*orf22*) was co-introduced, while excluding the adjacent TetR-family regulator that may negatively regulate cluster expression (Figures 1B, S3B).^28, 31^ The resulting recombinant strain produced BLA at titers over 10-fold higher than the native strain and maintained stable yields for at least seven days of fermentation (Figures 1C, 1D). This engineered strain overcame the BLA supply bottleneck, enabling comprehensive investigations of its structural, bioactivity, and mechanistic properties.

### Butyrolactol A synergizes with caspofungin in *Cryptococcus spp.* and resistant *Candida auris*

We initially identified BLA through its ability to potentiate caspofungin (CAP) activity against intrinsically resistant *Cryptococcus* at a single concentration in the screen. To further evaluate the breadth and mechanistic relevance of this synergy, we systematically assessed the dose-dependent antifungal activity of BLA alone and in combination with caspofungin across a panel of WHO-designated critical priority fungal pathogens. BLA alone exhibited moderate antifungal activity, with single-digit MICs against a range of yeast pathogens, including *C. neoformans*, *C. albicans*, *C. gattii*—an outbreak-associated species affecting immunocompetent individuals in North America—as well as the filamentous fungi *Aspergillus fumigatus* and *Trichophyton rubrum* (Table S5).

Notably, BLA displayed conserved synergy with caspofungin against *Cryptococcus* strains and *C. auris*, as indicated by fractional inhibitory concentration index (FICI) values < 0.5, reflecting robust combinatorial efficacy (Figure 2A, Table S6).^25^ This synergy significantly reduced CAP MICs, lowering them to or below the tentative CDC susceptibility threshold of ≤ 2 μg/mL for *C. auris*, when BLA was used at half its MIC (Table S6).^25, 32^ Although no established breakpoint exists for *C. neoformans*, this threshold serves as a useful benchmark for assessing enhanced susceptibility. The potentiating effect of BLA extended beyond CAP, as it also strongly synergized with the echinocandins micafungin and anidulafungin against *C. neoformans* H99 (FICI < 0.19), neither of which exhibited activity at concentrations up to 64 μg/mL (Figure S4A), further highlighting BLA’s role in restoring echinocandin efficacy. At sub-inhibitory concentrations (¼ MIC), the BLA–CAP combination achieved rapid and robust fungicidal activity, leading to complete clearance of *C. neoformans* cells with no detectable regrowth (Figure 2B). A similar effect was observed in isolates of multidrug-resistant *C. auris*—a WHO-designated critical priority pathogen that exhibits resistance to nearly all approved antifungal drugs, including both BLA and CAP when used individually.^33, 34^ Remarkably, the combination of 2 μg/mL BLA and CAP was the only tested treatment to fully eradicate fungal cells, maintaining sterilization for at least 15 days post-treatment (Figures 2B, S4B), underscoring the superior potency and durability of this combination strategy against highly refractory fungal pathogens.

**Figure 2.**
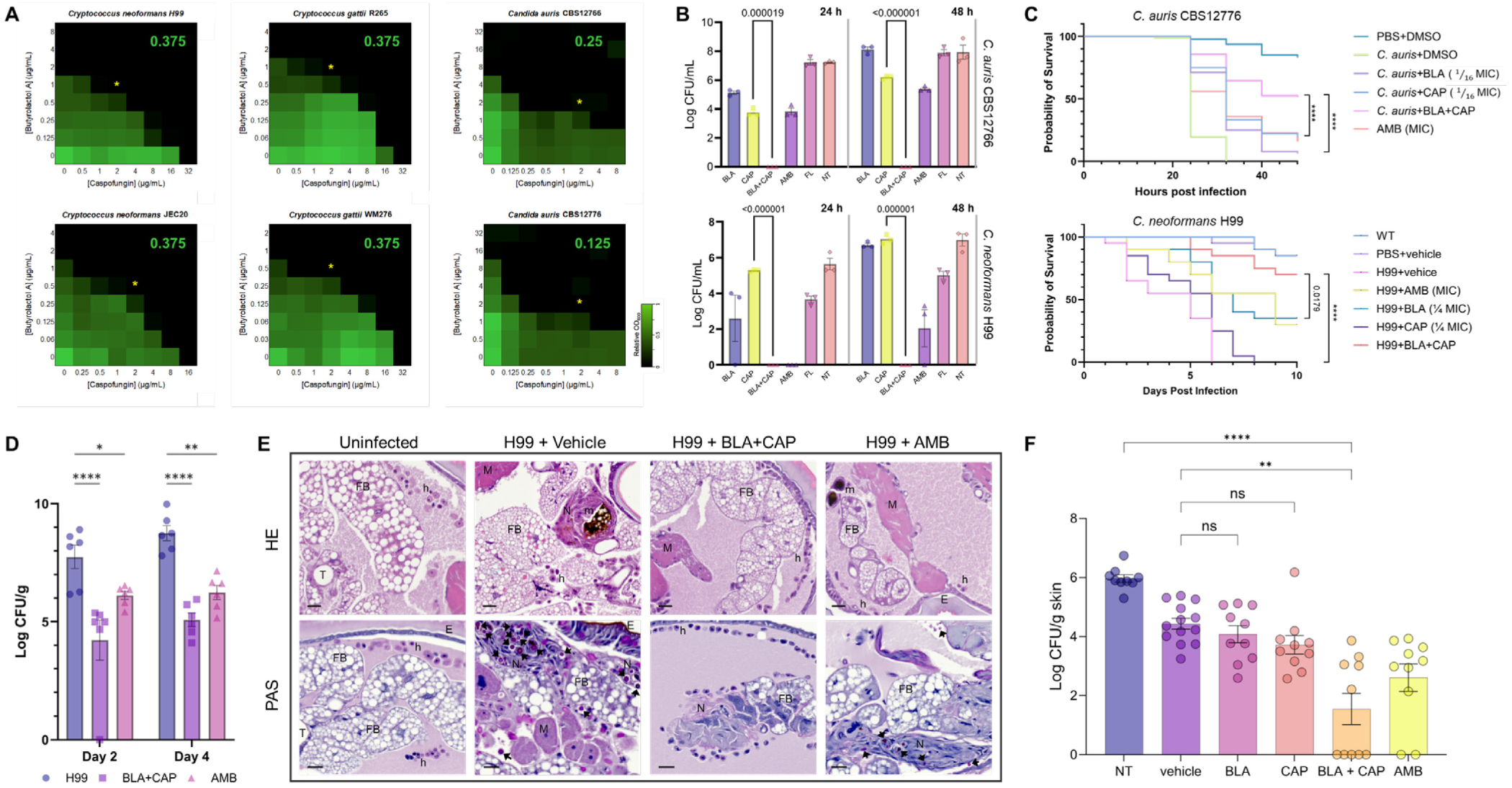
Butyrolactol A synergizes with caspofungin in *Cryptococcus* spp. and multidrug-resistant *Candida auris.* **(A)** Checkerboard assays depicted as heatmaps show the average growth of biological duplicates, normalized to no compound controls. Compound potentiation was assessed against *C. neoformans* H99 and JEC20, *C.s gattii* R265 and WM276, and *C. auris* CBS12766 and CBS12776. Relative growth is quantitatively represented by color (see scale bar at the bottom right). Fractional Inhibitory Concentration Index (FICI) values were calculated as described in the methods and are displayed in the top right corner of each checkerboard. FICI values below 0.5 indicate synergistic interactions. The yellow asterisk highlights the rescue concentration of butyrolactol A (BLA), which reduces the minimum inhibitory concentration (MIC) of caspofungin to 2 µg/mL. See also Figure S4A. **(B)** Fungicidal activity of BLA + CAP combination treatment. Fungal colony-forming units (CFUs) were quantified following 24 h and 48 h treatments with vehicle control (NT), butyrolactol A (BLA), caspofungin (CAP), their combination (BLA + CAP), amphotericin B (AmB), and fluconazole (FL). **(Top)** *C. auris* CBS12766: BLA (2 µg/mL), CAP (2 µg/mL), AmB (4 µg/mL), FL (64 µg/mL). **(Bottom)** *C. neoformans* H99: BLA (0.5 µg/mL), CAP (4 µg/mL), AmB (0.5 µg/mL), FL (8 µg/mL). Data are presented as mean ± SEM from three independent biological replicates. *p*-values were calculated using unpaired *t*-tests, statistically significant comparisons are indicated on the graph. Samples with no detectable CFUs (CFU = 0) were considered below the detection limit and are plotted as log₁₀ CFU/mL = 0 for visualization only. See also Figure S4B. **(C)** *C. elegans* were infected with *C. auris* CBS 12766 (upper) and *G. mellonella* were infected with *C. neoformans* H99 (bottom). Survival was monitored in the presence of butyrolactol A (BLA), caspofungin (CAP), the combination of both, amphotericin B (AMB)or without treatment. For each condition, 20 *C. elegans* worms were used, and survival was tracked over 48 hours across 3 independent trials. Similarly, 10 *G. mellonella* larvae were used for each condition, and survival was tracked over ten days after injection with ∼10^6^ per larvae in 3 independent trials. Survival was plotted using a Kaplan-Meier survival curve. Significance was determined using the Log-rank (Mantel-Cox) test, comparing monotherapy of CAP and BLA to combination therapy, with the corresponding p-values indicated as numerical values or **** for p < 0.0001. **(D)** *G. mellonella* larvae infected with *C. neoformans* H99 were treated with butyrolactol A (BLA) + caspofungin (CAP) at ¼ MIC each, amphotericin B (AMB, MIC), or vehicle as control (H99). Fungal burden (CFU per larva) was quantified at Day 2 and Day 4 post-infection. Data are presented as mean ± SEM from 6 larvae per group. Statistical significance was determined using Dunnett’s multiple comparisons test relative to the infected control group: *p < 0.05, **p < 0.01, ***p < 0.001, \*\*\*\**p < 0.0001.* **(E)** Histopathological analysis of *G. mellonella* tissues following treatment of *C. neoformans* infection. Representative histological sections of *G. mellonella* larvae harvested 2 days post-infection with *C. neoformans* H99, subjected to different treatments. Tissues were stained with hematoxylin and eosin (H&E) or periodic acid–Schiff (PAS) and imaged at 20× magnification. The panels show fungal morphology, host immune responses, and tissue integrity across treatment conditions. E, epithelial layer; FB, fat body; h, hemocyte; N, hemocyte nodule; M, muscle; m, melanin; T, trachea. Black arrows indicate encapsulated yeast cells of *C. neoformans* localized within hemocyte nodules, consistent with immune activation. Scale bar = 20 μm. See also Figure S4C. **(F)** Butyrolactol A and caspofungin combination reduces *C. auris* fungal burden in a murine skin colonization model. C57BL/6N mice were topically infected with *C. auris* CBS12776 (1 × 10⁸ CFU) and treated with vehicle, butyrolactol A (BLA, 10 mg/kg), caspofungin (CAP, 10 mg/kg), amphotericin B (AmB, 25 mg/kg), or BLA–CAP combination at subtherapeutic doses. Treatments were administered at 24, 26, 28, and 44 h post-infection. Fungal burden was quantified at 48 h from homogenized skin tissue. Each dot represents one animal, and data are presented as mean ± s.e.m. Statistical analysis was performed using the Kruskal–Wallis test followed by Dunn’s multiple comparisons test; *p < 0.05, **p < 0.01, ***p < 0.001, \*\*\*\**p < 0.0001.* See also Figure S4D.

The *in vivo* efficacy of this combination was further validated in *Caenorhabditis elegans* and *Galleria mellonella* infection models, which are widely used for studying fungal–host interactions due to their conserved innate immunity and experimental tractability.^35, 36^ In the *C. elegans*-*C. auris* infection model, low concentrations of BLA and caspofungin alone (1/16 MIC) had minimal effect on animal survival. In contrast, their combination significantly improved host survival compared to either monotherapy (Figure 2C), supporting the combinatorial efficacy of this approach against multidrug-resistant *C. auris in vivo*. A similar trend was observed in *G. mellonella* larvae infected with *C. neoformans* H99, where the combination treatment significantly prolonged survival relative to either agent alone at sub-therapeutic doses, with a survival probability of ∼70% at Day 10, markedly higher than other treatment groups (Figure 2C). Quantification of fungal burden in infected larvae revealed a rapid and pronounced reduction in viable *C. neoformans* cells as early as Day 2 post-infection in the BLA and CAP combination treatment group, which was sustained through Day 4 (Figure 2D), supporting a rapid and potent *in vivo* antifungal effect of the BLA–CAP combination, in line with our *in vitro* observations.

To further elucidate host–pathogen interactions and treatment effects at the tissue level, histopathological analyses were performed on *G. mellonella* larvae 48 h post-infection with *C. neoformans* H99 under different therapeutic regimens. In vehicle-treated controls, periodic acid– Schiff (PAS) staining revealed extensive fungal dissemination, with abundant yeast cells embedded within hemocyte nodules and diffusely infiltrating the fat body, digestive tract, and respiratory tissues (Figures 2E, S4C). Hematoxylin–eosin (H&E) staining showed a robust immune response characterized by widespread infiltration of histiocytic spindle-shaped hemocytes—reflecting activated immune cells and systemic inflammatory stress—into major organs, particularly the fat body and peritracheal regions.^37^ This was accompanied by a notable reduction in immature, unactivated hemocytes in subcuticular regions, reflecting systemic immune mobilization (Figures 2E, S4C). In contrast, combination treatment with BLA and CAP markedly reduced fungal burden and altered the histological landscape. Intact yeast cells were rarely detected, and hemocyte nodules containing fungal elements and melanin deposits were significantly smaller and less frequent, particularly in hemolymph-rich areas, reflecting effective fungal clearance and alleviation of host immune stress (Figures 2E, S4C).

The therapeutic potential of the BLA–CAP combination in a mammalian host was further assessed using a murine skin colonization model—a clinically relevant system that recapitulates superficial fungal persistence and transmission risk.^38^ *C. auris,* a highly multidrug-resistant pathogen, readily colonizes skin and serves as a reservoir for nosocomial outbreaks. Topical co-administration of BLA and caspofungin at subtherapeutic doses significantly reduced fungal burden, whereas monotherapies, including amphotericin B (25 mg/mL), showed no significant benefit over vehicle treatment (Figure 2F). Notably, mice receiving combination therapy exhibited no signs of morbidity or weight loss, indicating good tolerability (Figure S4D). These results extend the *in vitro* and invertebrate synergy of the BLA–CAP combination to a mammalian context and highlight its translational potential for treating clinically recalcitrant fungal infections.

Given the potent therapeutic activity of BLA, we next evaluated its cytotoxicity across multiple human cell lines to assess host selectivity. Using both ATP-based viability and LDH-release assays, BLA showed substantially lower cytotoxicity than amphotericin B in HEK293 (kidney), HepG2 (liver), and THP-1 (immune) cells (Table S7). In HEK293, the IC₅₀ values of BLA were 286 μg/mL (ATP) and 127 μg/mL (LDH), compared to 30.6 μg/mL and 9.4 μg/mL for amphotericin B. Similar trends were observed in HepG2 cells, while THP-1 cells were more sensitive to both compounds. Notably, BLA induced minimal LDH release at antifungal concentrations, indicating a low membrane-disruptive potential. These results support a favorable safety margin for BLA and underscore its selectivity for fungal over mammalian cells.

### Butyrolactol A perturbs sterol homeostasis without directly targeting ergosterol or the cell wall

Amphotericin B (AMB) exerts its fungicidal activity by binding and extracting ergosterol from lipid bilayers, a mechanism that also disrupts membrane integrity and contributes to its well-known cytotoxicity in human cells (Figure 3A). The AMB-resistant *C. albicans* strain ATCC 200955, which harbors a defective sterol biosynthesis pathway (Figures 3B–3C)^39^ is also resistant to other ergosterol-targeting agents such as fluconazole and terbinafine, as well as to BLA, showing a >64-fold increase in MIC compared to reference strains (Figure 3A, Table S5). This cross-resistance and related structures (i.e., polyketide polyols) between BLA and AMB (Figure 3D) suggested a shared mechanism of action.^40, 41^ However, supplementation with exogenous ergosterol, which dose-dependently reversed fungicidal activity of AMB, failed to rescue cells from BLA-induced killing (Figure 3E), indicating a mechanism distinct from ergosterol binding or sequestration.

**Figure 3.**
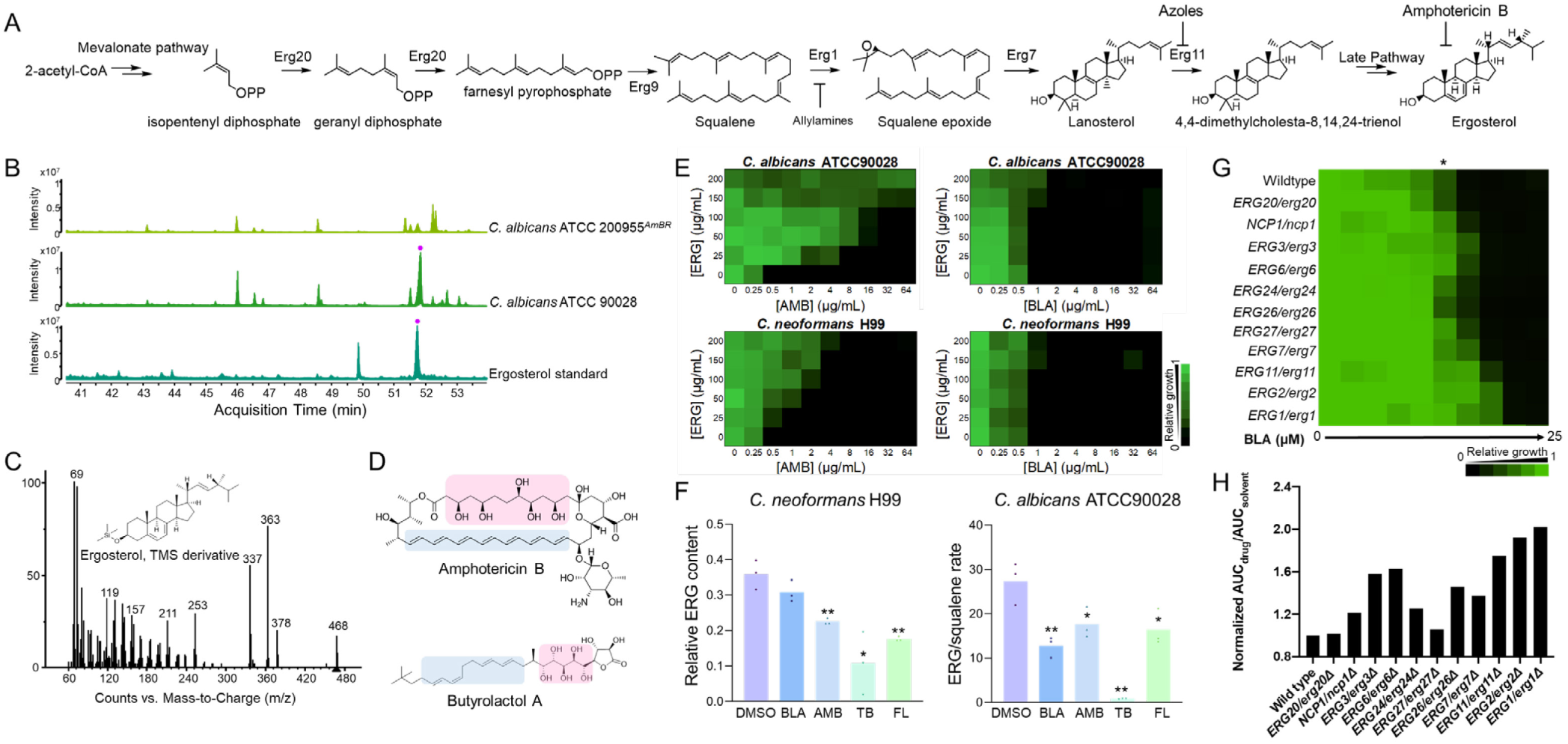
Butyrolactol A perturbs sterol homeostasis without directly targeting ergosterol or the cell wall. **(A)** Simplified schematic of the ergosterol biosynthesis pathway, along with antifungals targeting ergosterol or its biosynthesis. **(B)** Gas chromatography-mass spectrometry (GC-MS) analysis of ergosterol in amphotericin B-resistant *C. albicans* ATCC200955 (upper) and amphotericin B-sensitive *C. albicans* ATCC90028 (middle). The ergosterol standard is shown as a reference (bottom). **(C)** Chemical structure and GC-MS spectrum of ergosterol-TMS derivative. **(D)** Structural comparison of butyrolactol A and amphotericin B, highlighting structural similarities. The hydrophobic polyene region is indicated in blue, while the hydrophilic polyol structure is in pink. **(E)** Rescue effect of exogenous ergosterol (ERG) on cell viability during treatment with butyrolactol A (BLA), with amphotericin B (AMB) used as a positive control. Two-fold serial dilutions of ergosterol were added to *C. neoformans* H99 and *C. albicans* ATCC90028 in the presence of AMB or BLA, respectively. Biological duplicates were averaged and normalized to no-drug and no-ergosterol controls. Relative growth is quantitatively represented by color (see scale bar at the bottom right). **(F)** Relative ergosterol (ERG) abundance in *C. neoformans* H99 and *C. albicans* ATCC90028, treated or untreated with butyrolactol A and different agents targeting ergosterol biosynthesis: amphotericin B (AMB), terbinafine (TB), and fluconazole (FL). The median value of three biological replicates is represented by the height of each bar, with individual experimental values indicated by dots. Significance was determined by one-way ANOVA with Dunnett’s multiple comparisons test, comparing each treatment to the untreated control (* p < 0.05, ** p < 0.01, t-test). **(G)** *In vitro* susceptibility of *C. albicans* wild-type (WT) and heterozygous-deletion mutants to BLA. Dose-response assays were performed in YPD medium, with growth measured by absorbance at 600 nm after 24 hours at 30°C. Optical densities were averaged from duplicate measurements and normalized to the growth of untreated WT strains. Results are depicted as in Figure 2A (see bar for scale). (* indicates concentrations selected for plotting the growth curve). **(H)** Relative growth of *C. albicans* strains at 1.56 μM BLA (indicated by * in Figure 3G). The area under the growth curve (AUC) for strains in the presence of BLA was normalized to their AUC in the absence of BLA to eliminate baseline growth discrepancies. Data are presented as the mean of technical replicates.

To examine whether BLA impacts sterol homeostasis, we quantified ergosterol levels in *C. neoformans* and *C. albicans* after treatments. As expected, fluconazole and terbinafine significantly reduced ergosterol in *C. neoformans* and *C. albicans* by targeting *ERG11* and *ERG1*, respectively (Figures 3A, 3F). BLA treatment resulted in a modest reduction of ergosterol in *C. neoformans* and a more substantial (61%) decrease in *C. albicans*, indicating perturbation of sterol homeostasis with species-specific differences in sensitivity (Figure 3F). Given the stronger response observed in *C. albicans*, we selected this species to further investigate whether BLA targets enzymes within the ergosterol biosynthesis pathway. Sensitivity profiling of eleven heterozygous *ERG* deletion mutants revealed no evidence of hyper-susceptibility to BLA, suggesting that the enzymes in ergosterol biosynthesis are unlikely to represent its direct molecular target (Figure 3G). Conversely, heterozygous deletion of most *erg* genes conferred resistance to BLA (Figure 3H), suggesting that membrane composition is a key modulator of susceptibility in *C. albicans*. Additionally, osmotic protection with sorbitol did not affect the fungicidal activity of BLA, in contrast to caspofungin, whose MIC increased by 64-fold under the same conditions (Table S5), indicating that BLA does not kill by compromising cell wall integrity.

### Identification of butyrolactol A as an inhibitor of the flippase complex Apt1–Cdc50

Although our genetic and phenotypic analyses suggested that BLA perturbs sterol homeostasis, the underlying molecular mechanism remained elusive. To identify potential targets, we selected for BLA resistance and then performed whole-genome sequencing.^42^ Two BLA-resistant *C. neoformans* H99 mutants were obtained through serial passaging in subinhibitory BLA concentrations, followed by final selection on 4×MIC BLA agar (Figure S5A). Whole-genome analysis revealed a shared mutated gene, *CNAG_06469* (*APT1*), which encodes a P4-ATPase, also known as lipid flippase, belonging to the Drs2-related P4A subfamily (Figure 4A).^43^ The *APT1* mutations included a 22-base tandem duplication insertion at nucleotide positions 557845-557867, causing a frameshift in the C-terminal tail (RA3), and G266V substitution within the actuator (A) domain (RA4) (Figure 4A, S5B). To determine if these mutations are responsible for BLA resistance, we performed genetic backcrosses with KN99a. Both RA3 × KN99a and RA4 × KN99a F1 progeny displayed nearly 1:1 segregation patterns of the resistance phenotype, consistent with inheritance of a single genetic locus (Figure 4B, Table S8). Genotyping confirmed that resistance co-segregated with the mutant *APT1* allele, demonstrating that these *APT1* mutant variants confer BLA resistance (Table S8).

**Figure 4.**
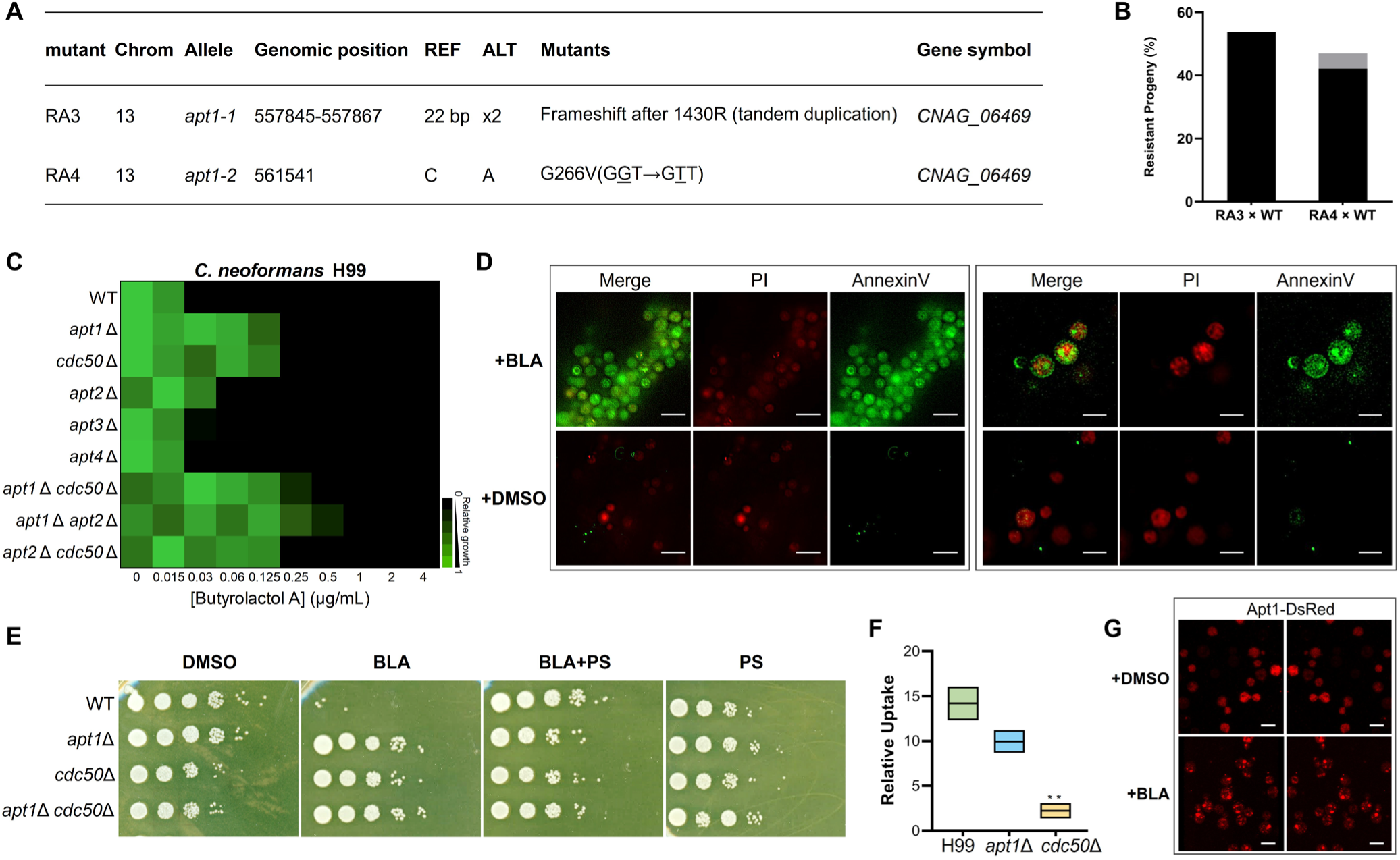
Identification of butyrolactol A as an inhibitor of the flippase complex Apt1– Cdc50. **(A)** Genotypes of BLA-resistant mutants, carrying mutations in *APT1* (*CNAG_06469*). See also BLA susceptibility results in Figure S5A and the mutations mapped onto the Apt1 structure in S5B. **(B)** Results of the genetic cross between BLA-resistant mutants and the wild-type strain (KN99**a**). The *y*-axis represents the percentage of resistant progeny, with the inherited resistant genotypes from the parent strains shown in black. The *x*-axis shows different cross tests between independent resistant mutants RA3 and RA4 (as mentioned in Figure 4A) and the wild-type strain KN99a (WT). See also Table S8. **(C)** Dose-response assays were performed against wild-type *C. neoformans* (H99) and various flippase-compromised mutants. Growth was measured using absorbance at 530 nm in SDB medium at 30 °C after 48 hours. Growth relative to untreated wild-type control wells is presented in a heatmap format (see scale bar). Each colored box represents the mean of duplicate experiments. **(D)** *C. neoformans* H99 cells in log phase were treated with BLA (upper) or an equivalent amount of DMSO (bottom) and were co-incubated with Annexin V Alexa Fluor™ 488 and Propidium Iodide (PI), then observed using fluorescence microscopy. The Annexin V Alexa Fluor™ 488 signal was detected in the FITC channel, and the PI signal was detected in the TexRed channel. Scale bars are shown in each image: 10 μm (left column) and 5 μm (right column). **(E)** The effect of exogenous phosphatidylserine (PS) on the viability of wild-type *C. neoformans* H99, *apt1*Δ, *cdc50*Δ, and *apt1*Δ *cdc50*Δ mutants in the presence or absence of BLA. Strains were inoculated in SDB medium treated with BLA, PS, or a combination of BLA and PS for 24 hours. Equivalent vehicle DMSO was used as a control. Ten-fold serial dilutions of the 24-hour cultures were spotted onto yeast extract peptone dextrose (YPD) plates. Plates were incubated at 30 °C for 2 days before imaging. **(F)** The relative intracellular concentrations of BLA accumulated in *C. neoformans* H99 and *apt1*Δ and *cdc50*Δ mutants were measured after 15 minutes of treatment. Data are presented as mean ± SD of biological triplicates. Statistical significance was determined using multiple unpaired t-tests with Holm-Sidak’s multiple comparisons correction, comparing all conditions to wild-type H99. p-values: ** <0.01. **(G)** Localization of DsRed-Apt1p in *C. neoformans* H99 cells with (upper row) and without (lower row) BLA treatment. Representative Airyscan super-resolution 3D images. Z-stacks were reconstructed into 3D projections. The left and right panels show the same cell from two different viewing angles to illustrate its spatial distribution. Scale bar, 5 μm.

Finally, the resistance of a cryptococcal *apt1*Δ mutant to BLA supported Apt1 as a likely target of BLA and suggested that the mutations identified in RA3 and RA4 represent loss-of-function alleles (Figure 4C). Most P4-ATPases function as binary complexes with a β-subunit from the CDC50 family. The only member of this family in *C. neoformans*, Cdc50, serves as a β-subunit for Apt1, forming a stable complex that functions as a lipid flippase.^44, 45^ Therefore, the susceptibility of *cdc50*Δ and *apt1*Δ *cdc50*Δ mutants was characterized. We observed that depletion of CDC50 resulted in severe growth defects in RPMI medium, which mimics *in vivo* physiological conditions, particularly at 37 °C. This finding aligns with previous reports of impaired survival of *cdc50Δ* mutants in mouse models, underscoring the critical role of CDC50 in the *in vivo* survival of fungal pathogens.^22, 44^ In acidic, carbon-enriched SDB medium, commonly used for rapid fungal culture, the *cdc50*Δ and *apt1*Δ *cdc50*Δ mutants displayed similar resistance to BLA as *apt1*Δ, suggesting that BLA targets the flippase complex (Figure 4C). Besides Apt1, three additional P4-ATPase genes were identified in the cryptococcal genome (*APT2*, *APT3*, and *APT4*). Only the *apt2*Δ mutant exhibited modest BLA resistance, likely due to its close phylogenetic relationship to Apt1 (Figures 4C, S5C). The enhanced resistance observed in the double mutant *apt1*Δ *apt2*Δ suggests partial functional redundancy between Apt1 and Apt2 (Figure 4C). Nevertheless, Apt1 is the principal target of BLA, underscoring its selective action among P4-ATPases.

Apt1 and Apt2 are closely related to the plasma membrane P4-ATPases Dnf1/2 from *S. cerevisiae*, which have broad substrate specificity and do not require a phosphate headgroup for lipid substrate recognition (Figure S5C).^46^ The transmembrane flippase, often associated with an indispensable β-subunit belonging to the Cdc50 family, drives an inward translocation of various phospholipids to establish an essential asymmetric lipid distribution of the membrane with phosphatidylserine (PS) and phosphatidylethanolamine (PE) concentrated within the cytosolic leaflet.^47, 48^ Cdc50 depletion results in a loss of membrane asymmetry that exposes PS on the extracellular leaflet.^22, 44^ The inhibitory effect of BLA on the flippase function of Apt1–Cdc50 was validated by monitoring PS accumulation on the outer leaflet using green fluorescent conjugated annexin V.^22^ The significantly elevated PS exposure upon BLA treatment supports a model where BLA inhibits the flippase complex, thereby blocking phospholipid translocation and consequent disruption of membrane asymmetry (Figure 4D).

The amphipathic structure of BLA resembles natural flippase substrates, featuring a hydrophobic tail and a hydrophilic headgroup that interacts with the protein.^46, 49^ Given the broad substrate specificity of the Apt1–Cdc50 complex, we reasoned that BLA might be mistakenly loaded as a substrate analog,^45^ inhibiting flippase activity in a competitive manner, thereby obstructing lipid entry and leading to PS accumulation on the extracellular leaflet.^48^ To test this competitive model, exogenous PS was supplied as a substrate competitor. While PS alone did not affect cryptococcal growth, it effectively restored the BLA-induced severe growth defect to the level of untreated/resistant cells (Figure 4E). Other phospholipids such as phosphatidylcholine (PC), phosphatidylethanolamine (PE), and the phospholipid precursor phosphatidic acid (PA) also conferred some rescue of *C. neoformans* growth, whereas triacylglycerol (TAG), which lacks a hydrophilic headgroup, did not (Figure S5D).

These results established flippase inhibition–mediated membrane remodeling as a distinct antifungal mechanism of BLA and prompted further investigation into the role of the flippase complex in BLA uptake.^50^ Deletion of *cdc50* significantly reduced BLA uptake, and *apt1* deletion moderately decreased its uptake, possibly due to the redundant contributions of the flippase catalytic subunits, suggesting an interaction between BLA and the Apt1–Cdc50 complex (Figures 4C and 4F). In contrast, both deletions resulted in increased uptake of AMB, which is consistent with the distinct MOA proposed for the two compounds (Figure S5E).^22, 43^ In addition to serving as an integral part of the lipid translocating machinery for P4-ATPases, Cdc50 also functions as a chaperone, facilitating the exit of flippases from the endoplasmic reticulum (ER) to their proper subcellular destinations.^51^ Under normal conditions, Apt1-DsRed primarily cycles between the Golgi ^52^ endosomes, and cell membrane, appearing as punctate structures distributed throughout the cell, as visualized by super-resolution Airyscan 3D imaging, which is consistent with previous reports.^53, 24^ Upon BLA treatment, Apt1-DsRed undergoes a marked redistribution, with loss of the typical punctate pattern and the formation of prominent peripheral clusters at the cell periphery (Figure 4G). This altered localization suggests that BLA disrupts normal intracellular trafficking and retrieval of Apt1, potentially through inhibition of Cdc50 function.^54, 55^

### Cryo-EM reveals butyrolactol A binding to Apt1–Cdc50 and obstruction of the lipid transport channel

Cryo-electron microscopy (cryo-EM) studies were conducted to further elucidate the molecular mechanism of BLA-mediated flippase inhibition and its interactions with Apt1–Cdc50. We heterologously co-expressed the full-length *C. neoformans* Apt1 and Cdc50 in *S. cerevisiae* BY4741 *pep4*Δ and purified the heterodimeric protein complex using the detergents lauryl maltose neopentyl glycol (LMNG) and cholesteryl hemisuccinate (CHS). ATP was supplemented during the purification wash steps to remove endogenously bound lipids from lipid-bound flippases.^56^

As a P4-ATPase, Apt1–Cdc50 operates via the Post-Albers mechanism, where cyclic phosphorylation and dephosphorylation in the cytoplasmic ATPase domain drive phospholipid translocation, generating intermediate states with distinct substrate affinities.^57^ In the E2P state, a hydrophilic cleft opens toward the extracellular/luminal leaflet, enabling substrate lipid loading and polar headgroup binding, representing a high-affinity state for transported substrates.^48^ Substrate lipids have been captured at the entry sites of their respective E2P-state flippases, serving as lipid cargos poised for translocation.^58, 59, 60, 61^ Recognizing that BLA competes with the substrate lipid for the same binding site, we locked Apt1–Cdc50 in the E2P state using the phosphate analog BeF_3_^−^ and resolved the BLA-bound cryo-EM structure at an overall resolution of 3.0 Å (Figures 5A and S6, Table S9). To compare and validate the competition model, the E1 state of the flippase complex was also determined (Figure S7, Table S9).

**Figure 5.**
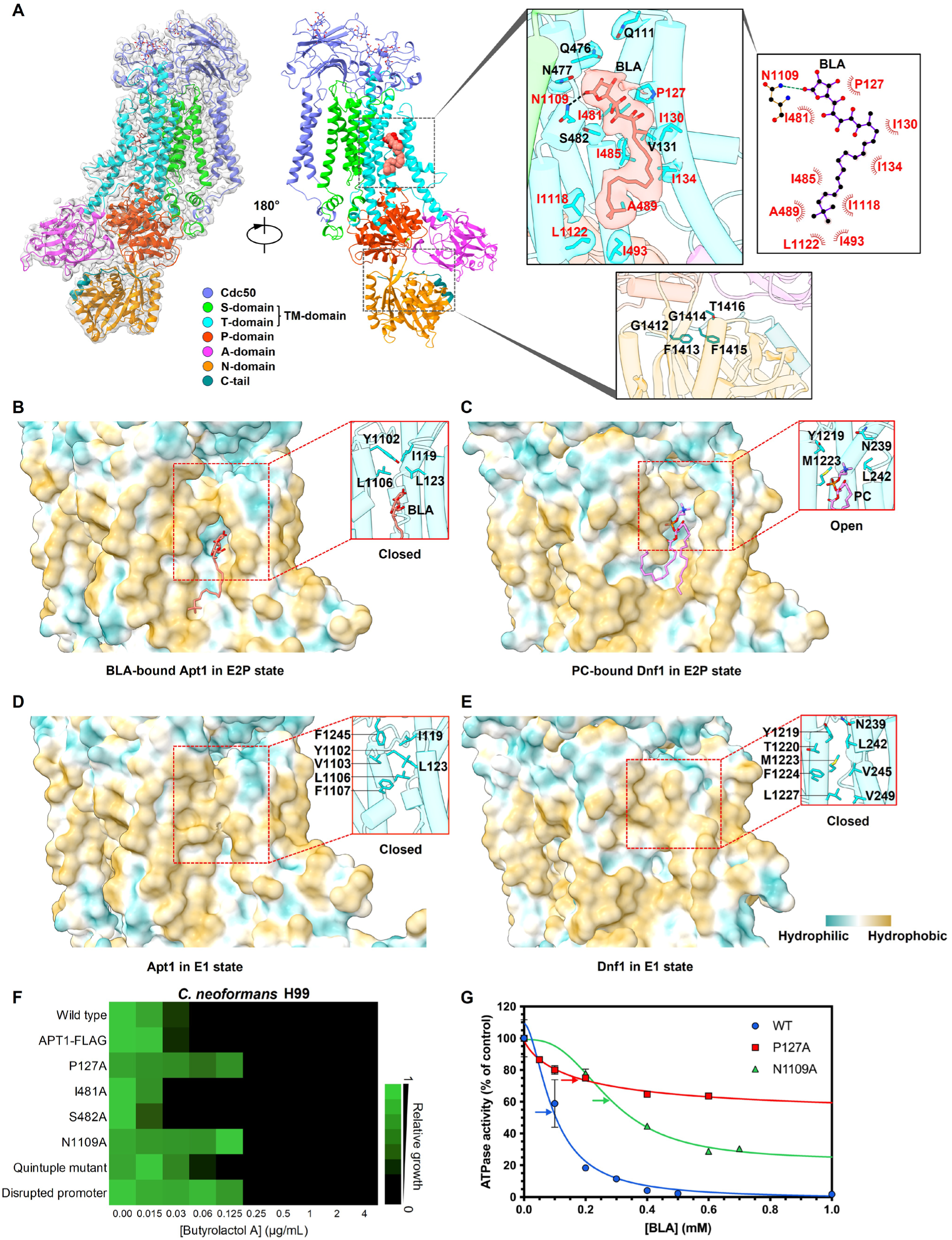
Cryo-EM reveals butyrolactol A binding to Apt1–Cdc50 and obstruction of the lipid transport channel. **(A)** Overall structure of *C. neoformans* Apt1–Cdc50 bound with BLA (salmon spheres) in the E2P state with major domains shown in different colors. The global view on the left is also shown with the cryo-EM map density fit. TM-domain, transmembrane domain; S-domain, support domain; T-domain, transport domain; P-domain, phosphorylation domain; A-domain, actuator domain; N-domain, nucleotide-binding domain. Top right: enlarged view of BLA binding site with the EM density of BLA and its surrounding residues superimposed on the atomic model and shown as transparent surfaces. BLA and residues surrounding the BLA binding site are shown in sticks. The key residues forming either H-bonds (indicated by dashed line with a cutoff distance of 3.5 Å) or hydrophobic interactions (cutoff distance: 4.0 Å) with BLA are labeled in red. A 2D plot of BLA and its interacting residues is also shown next to the corresponding 3D view. Bottom right: enlarged view of the autoregulatory C-tail with the GFGFT motif shown in teal sticks. **(B)** Binding of BLA to Apt1 in the E2P state induces closure of the lipid entry site shown in hydrophobicity surface. BLA is shown in salmon sticks and a cartoon representation of the lipid entry site boxed in red is also shown in the inset with the entry closing residues in cyan sticks. **(C)** Lipid entry site of *S. cerevisiae* Dnf1 in the E2P state remains open upon binding of the substrate lipid PC as shown in the hydrophobicity surface (PDB ID 7KYC). PC is shown in magenta sticks and a cartoon representation of the lipid entry site in the open state boxed in red is also shown in the inset. **(D)** Lipid entry site of Apt1 in the E1 state is closed as shown in the hydrophobicity surface (PDB ID 9DZV, this study). A cartoon representation of the closed lipid entry site boxed in red is also shown in the inset with interacting residues in cyan sticks. **(E)** The closed lipid entry site of *S. cerevisiae* Dnf1 in the E1 state as shown in the hydrophobicity surface (PDB ID 7KY6). A cartoon representation of the entry closing boxed in red is also shown in the inset with interacting residues in cyan sticks. **(F)** BLA susceptibility of wild-type *C. neoformans* H99 and various mutants, including FLAG-tagged Apt1 (Apt1-FLAG), single substitutions P127A, I481A, S482A, N1109A, a quintuple mutant (I130A, I134A, I485A, I493A, I1118A), and the Apt1-FLAG promoter disrupted mutant. Relative growth is shown in heat-map format (see scale bar). Each colored box represents the mean of duplicate measurements. Refer to the transcriptional and translational levels of Apt1 in wild-type and mutant strains in Figures S9B and S9C. **(G)** BLA inhibition of PS-stimulated ATPase activity of wild-type *C. neoformans* Apt1 and mutants P127A and N1109A in the presence of 0.05 mM PS and increasing concentrations of BLA. The control activity in the absence of BLA was taken as 100% for each protein and data were fitted to an inhibitory dose-response equation with variable slope and IC_50_ indicated by the arrow. Data points represent the mean ± SD in triplicate (n = 3) for WT and duplicate (n = 2) for mutants. Refer to Figure S9D for activity comparison between WT and mutants without BLA.

An elongated EM density was observed within the lipid entry groove, aligning closely with the BLA molecule. The structure is clearly in the expected E2P state, supported by the well-ordered autoregulatory C-tail composed of the GFGFT motif and a short α-helix (Figure 5A).^57, 62^ Notably, the BLA binding prominently induced the closure of the lipid entry site (Figures 5A, 5B). The γ-butyrolactone moiety of BLA is stabilized by hydrophobic interactions with I481 from the PISL motif of TM4 (Transmembrane Helix 4) and P127 from TM2 (Figures 5A, 3D), while the hydroxylated γ-lactone ring is positioned just over 3.5 Å from the conserved residues Q111, Q476, and N477, with potential water-bridged hydrogen bond interactions. The carbonyl group of the lactone ring forms a hydrogen bond with N1109 of TM6, anchoring the lipid-like BLA headgroup to the substrate access channel (Figures 5A, 3D). Additionally, two pairs of hydrophobic residues (I119 and Y1102, L123 and L1106) seal the entry site of the substrate channel, preventing further substrate lipids from entering from the membrane matrix (Figure 5B). This binding mode contrasts with that of *S. cerevisiae* Dnf1 in the same E2P state, where the channel entry remains fully open when bound to a substrate lipid such as PC (Figure 5C).^59^ Interestingly, the aliphatic tail of BLA appears to jam the transport channel by adopting a kinked configuration by virtue of its conjugated double bonds. The BLA tail extensively interacts with the so-called ’hydrophobic barrier’ residues, including I130, I134, I485, A489, I493, I1118, and L1122 (Figures 5A, 5B, 3D).^48^ Despite the distinct effects of lipid-like BLA and substrate lipid PC on the conformation of the lipid transport channel (Figures 5B, 5C, S8A), the E1 states of Apt1 and Dnf1 are similar as both feature a closed channel that is incompatible with substrate binding at the entry site (Figures 5D, 5E, S8B).^59^ Collectively, these findings suggest a BLA inhibition mechanism wherein BLA binding induces closure of the lipid entry site and obstructs the transport channel with its kinked tetraene tail, effectively stalling the flippase in an abnormal E2P-like state.

Mutations of the key residues involved in BLA binding (Figure 5A) were prepared using a TRACE (Transient CRISPR-Cas9 coupled with Electroporation) strategy, which overcomes the challenges of genetic manipulation in *Cryptococcus* spp^63^. This approach resulted in 6% correctly integrated constructs, with an epitope FLAG-tag added to the C-terminus of wild-type and mutant Apt1 simultaneously, as verified by sequencing (Figure S9A). The mutations introduced in *APT1* did not significantly affect the flippase’s transcriptional or translational expression levels (Figures S9B-S9C). Subsequent susceptibility tests identified residues P127 and N1109 as crucial for BLA binding, as single mutations at either site significantly increased BLA resistance, mirroring that seen in the *apt1* promoter-disrupted mutant in which binding of BLA to the flippase is presumably abolished (Figure 5F). Interestingly, while both alanine substitutions at the equivalent site in the PS flippase ATP11C (i.e., P94A and N912A) exhibited comparable PS-dependent ATPase activities,^56^ the P127A and N1109A Apt1 mutants lost over 94% PS-dependent ATPase activities compared to the wild-type protein (Figure S9D). Crucially, the alanine substitutions of N1109 and P127 increased IC_50_ of the inhibitor BLA by 3-fold and 1.9-fold, respectively (Figure 5G), corroborating that BLA competes with PS for cargo binding in the cryptococcal flippase Apt1– Cdc50. In contrast, single mutations I481A and S482A within the conserved PISL substrate-binding motif did not confer BLA resistance. Given that the equivalent mutation of I481A in ATP11C (i.e., V357A) reduces PS-dependent ATPase activity by over 80% compared to the wild-type enzyme^56^ and the equivalent of S482 in ATP8B1 (i.e., S403) has been implicated in substrate recognition,^58^ we propose that I481 and S482 preferentially bind natural substrate lipids such as PS over the inhibitor BLA. This reduced natural cargo binding in these mutants may increase BLA binding, thereby mitigating resistance. Indeed, the S482A substitution increased BLA sensitivity two-fold, supporting the crucial role of S482 in stabilizing natural cargos, thereby enhancing the competitive binding advantage of BLA and rendering the mutant more susceptible (Figure 5F). Besides these conserved residues in anchoring the BLA headgroup, the middle “hydrophobic barrier” region extensively interacted with BLA’s uniquely folded tetraene tail, a feature rarely seen with natural cargos (Figures 5B, 5C). The quintuple mutant, bearing I130A, I134A, I485A, I493A, and I1118A substitutions within the hydrophobic barrier, showed a consistent 2-fold increase in MIC (Figure 5F), suggesting that the tail typically helps stabilize itself within the transport channel, a function that the mutant residues can no longer effectively perform.

### BLA inhibition of Apt1–Cdc50 disrupts membrane architecture and trafficking

BLA traps the Apt1–Cdc50 complex in a dead-end state, preventing the flippase from completing the catalytic cycle. Previous studies have highlighted the overlapping functions of Apt1 and Cdc50 in maintaining membrane integrity,^44, 45^ which may account for the cell death observed following BLA-mediated inhibition. Notably, BLA treatment caused dose-dependent ATP leakage, with intracellular ATP levels dropping 13-fold at 3 μM and 38-fold at 6 μM BLA, respectively, alongside a corresponding rise in extracellular ATP (Figure 6A). Membrane disruption became evident within 30 minutes and intensified over time, as indicated by a significant increase in ATP leakage (Figure 6B). These findings suggest that BLA binding to the flippase complex compromises membrane integrity, leading to substantial leakage of cellular contents and ultimately cell death (Figure S10A).

**Figure 6.**
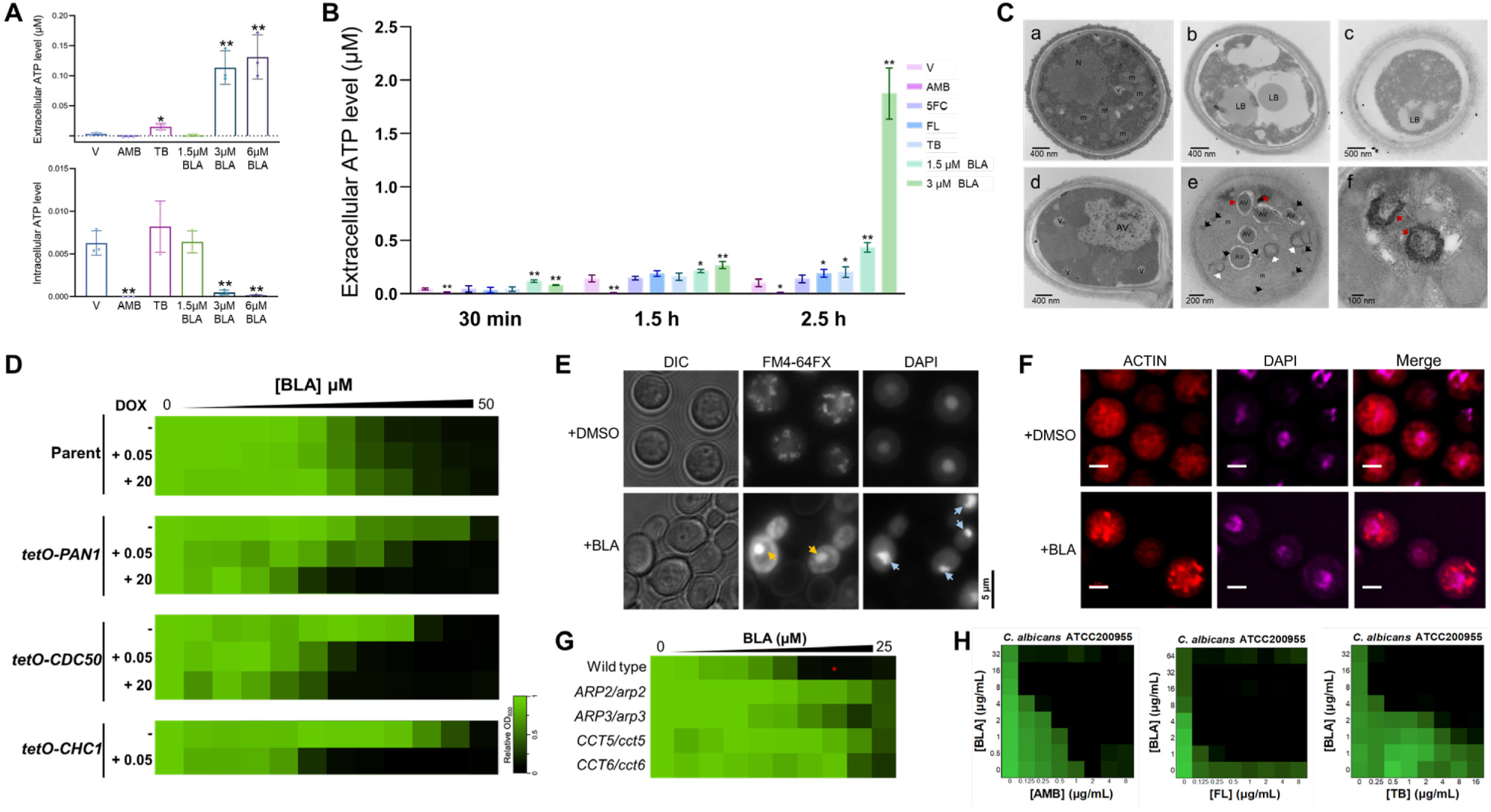
BLA inhibition of Apt1–Cdc50 disrupts membrane architecture and trafficking. **(A)** Cellular and extracellular ATP levels were measured following treatment. Log-phase *C. neoformans* H99 cells were cultured in SDB medium in the presence of amphotericin B (AMB), terbinafine (TB), butyrolactol A (BLA), or vehicle (-). Data are expressed as the mean ± SD of biological triplicates. Statistical significance was determined by multiple *t*-tests compared to the vehicle control (\**P* < 0.05, \*\**P* < 0.01). **(B)** ATP leakage was tracked by the extracellular ATP of *C. neoformans* H99 measured at 0.5-, 1.5-, and 2.5-hours post-treatment. 5-fluorocytosine (5FC), Fluconazole (FL). Experiments were performed in triplicate, and error bars represent standard deviations (n = 3). Statistical significance was assessed by multiple *t*-tests, with significance denoted by asterisks (\**P* < 0.05, \*\**P* < 0.01) for each treatment compared to the vehicle control. **(C)** *C. neoformans* accumulates abnormal membrane structures following BLA treatment. Cells were grown in SDB medium with BLA or vehicle, fixed, and visualized using transmission electron microscopy (TEM). Vehicle (DMSO) treated cells are shown in (a), while cells treated with 1 µM BLA are displayed in (b-d), and cells treated with 0.5 µM BLA in (e-f), with (f) showing a magnified view. Membrane defects in BLA-treated cells include enlarged atypical vacuoles (AV), double-membrane rings, crescent-shaped structures (white arrowheads), accumulated vesicles (black arrowheads), electron-dense misfolded membranes (red arrowheads), lipid bodies (LB), nuclei (N), mitochondria (M), and vacuoles (V). **(D)** BLA two-fold dose-response assays were conducted using wild-type and conditional expression *C. albicans* strains of *PAN1* (tetO-*PAN1/pan1Δ), CDC50* (tetO-*CDC50/cdc50Δ*), and *CHC1* (tetO-*CHC1/chc1*Δ) with varying concentrations of doxycycline (DOX, 0.05 µg/mL and 20 µg/mL). Growth was measured by absorbance at 600 nm after 48 hours at 30°C. Data are presented as a heatmap, with color representing relative growth, normalized to no-drug controls. **(E)** Endocytosis was assessed in *C. neoformans* H99 treated with or without BLA. Mid-log-phase cells were stained with FM 4-64 and DAPI, then visualized by fluorescence microscopy (100×). Yellow arrowheads indicate abnormal vacuoles/endosomes, and blue arrowheads indicate DAPI-stained nuclei. Scale bar (5 µm) is located at the bottom left. **(F)** The effect of BLA on actin cytoskeleton organization in *C. neoformans* H99 was examined by epifluorescence microscopy. Early-log-phase cells treated with low concentrations of BLA, or vehicle were fixed and stained with Alexa Fluor 568 phalloidin, and mounted with DAPI. Scale bar = 2 µm. **(G)** *In vitro* BLA susceptibility of *C. albicans* wild-type (WT) and heterozygous deletion mutants for genes involved in actin assembly. Dose-response assays were performed in YPD medium, with growth measured by absorbance at 600 nm after 24 hours at 30 °C. Data are depicted as in panel (D). (* indicates concentrations selected for growth curve plotting in Figure S10D). **(H)** Synergy plots for BLA in combination with drugs targeting ergosterol or its biosynthesis. Checkerboard assays, presented as heatmaps, show the average growth inhibition of *C. albicans* ATCC200955 across biological triplicates, normalized to no-compound controls. Fractional Inhibitory Concentration Index (FICI) values are shown in the top right of each heatmap, with values <0.5 indicating synergistic interactions.

The fungicidal activity of BLA relies on effective binding to the functional Apt1-Cdc50 complex, as mutations in Apt1 or the deletion of either component confer resistance to BLA. Consistently, a nonsense mutation in *CNAG_03870*, encoding a Δ8-fatty acid desaturase essential for glucosylceramide (GlcCer) biosynthesis,^64, 65^ conferred over 100-fold resistance to BLA only in the absence of APT1, suggesting that BLA activity requires both protein target and a specific lipid environment (Figure S10B). Since Apt1–Cdc50 mediates GlcCer transport and membrane architecture,^45^ this lipid-dependent mechanism of action reveals a previously unrecognized synthetic route to resistance, underscoring the complexity and selectivity of BLA’s antifungal activity. Conversely, galactose-induced overexpression of the Apt1–Cdc50 in *S. cerevisiae* resulted in pronounced hypersensitivity to BLA, suggesting that the stabilized interaction between BLA and Apt1–Cdc5 is toxic to the cell (Figure S10C). This mechanism resembles that of the topoisomerase I (Top1) inhibitor camptothecin, which does not directly damage DNA but rather traps Top1 in the DNA cleavage complex conformation. The resulting Top1 protein-DNA covalent adducts interfere with DNA replication and transcription, resulting in cytotoxicity.^66^

Transmission electron microscopy (TEM) provided direct visual evidence of membrane disruption by BLA. Untreated cells displayed a typical ultrastructural aspect of cryptococcal morphology (Figure 6C, a), while treatment with 1 μM BLA resulted in significant membrane detachment from the cell wall, indicative of impaired membrane architecture (Figure 6C, b-d). Additionally, abnormal cytoplasmic vacuoles containing lipid bodies were observed, along with large vacuole-like structures over 1 μm in diameter containing fragmented endomembrane pieces (Figure 6C, b-d). These structures resembled the giant vacuoles seen in *apt1*Δ mutants, resulting from the disrupted vacuolar organization.^67^ To investigate the disruption process, the BLA concentration was reduced to 0.5 μM, revealing extensive distorted membranous structures, including electron-dense, stacked membranes (red arrowhead) and vesicles (black arrowhead), enlarged atypical vacuoles (AV), mitochondrial swelling, and Berkeley body-like double-membrane compartments (∼200 nm), which are widely seen in secretory impaired mutants such as *sec7*, *sec14*, and *drs2* (Figure 6C, e-f).^68, 69^ BLA’s inhibition of the exocytosis pathway was further confirmed in conditionally-repressible *C. albicans* GRACE mutants.^70^ Transcriptional repression of clathrin heavy chain gene (*CHC1*), *CDC50*, and *PAN1* conferred hypersensitivity to BLA, while overexpression of the transcripts resulted in resistance in *C. albicans* (Figure 6D).^52, 71^ Pan1, a homologue of Esp15, localizes to clathrin-coated pits and functions in the same or parallel pathway as Chc1 and Cdc50 for clathrin-coated vesicle formation, suggesting that BLA inhibits the clathrin-dependent secretory pathway.^68, 72^

Given the critical roles of membrane organization in trafficking, the BLA effect on endocytosis was further investigated by assessing the uptake of lipophilic styryl dye FM4-64, which intercalates into the plasma membrane and is internalized via the endocytic pathway, labelling endosomes, vacuoles, and vesicle cycling, as observed in untreated *C. neoformans* cells (Figure 6E).^73^ Intensely fluorescent large formations accumulated in BLA-treated cells along with persistent signal retained on the cell membrane, indicating that BLA blocks endocytosis and phospholipid cycling (Figure 6E).^68^ Additionally, BLA induced significant morphological changes, suggesting disruption of the actin cytoskeleton. The high-affinity F-actin probe, phalloidin, conjugated to Alexa Fluor 568 dye revealed an aberrant actin distribution in *C. neoformans* cells treated with a low concentration of BLA. Unpolarized actin clumps (∼0.8 µm diameter) were observed in ∼32% of treated cells, a phenotype typically associated with yeast mutants lacking actin-regulating kinase activity, likely due to unregulated actin nucleation complex Arp2/3 (Figures 6F, S10D).^74^ This suggests that BLA impaired actin motility and polarization likely through interaction with Apt1, which is functionally linked to Drs2 and involved in actin polarization and endocytosis.^43, 75^ In addition, heterozygous deletion of the actin assembly genes *ARP2*, *ARP3*, *CCT5*, or *CCT6* conferred strong resistance to BLA in *C. albicans* (Figure 6G), further indicating BLA tends to impair actin- and clathrin-mediated endocytosis while sparing the alternative Arp2/3-independent endocytic pathway (AIE).^76^

In addition to influencing actin cytoskeletal organization, membrane lipid composition is essential for determining sensitivity to drugs targeting ergosterol biosynthesis,^75^ such that flippase-deficient mutants *cdc50*Δ and *apt1*Δ exhibited increased susceptibility to fluconazole (Figure S10E).^22, 43^ The combination of BLA with amphotericin B, fluconazole, or terbinafine was investigated, revealing a significant synergistic interaction between these agents (Figure 6H). Though P4 ATPases are widely found in eukaryotic cells and are essential for cell membrane homeostasis, BLA showed much lower cytotoxic and hemolytic toxicity than amphotericin B (Figure S10F, Table S7). The high selectivity of BLA may be attributed to the redundant roles of flippases in mammals (Figure S5C).

### Butyrolactol A potentiates the fungicidal activity of caspofungin against *C. neoformans* by inhibiting the Apt1–Cdc50 complex

We propose a model in which BLA competitively inhibits the Apt1**–**Cdc50 complex, disrupting membrane lipid asymmetry and consequently leading to impaired membrane architecture and the blockage of clathrin-dependent membrane trafficking (Figure 7A). A recent study revealed that Cdc50 is essential for the innate echinocandin resistance of *Cryptococcus* spp. by preventing the uptake of the drug and maintaining intracellular calcium homeostasis that regulates cell wall synthesis and stress responses.^22, 23^ Both *cdc50*Δ and *apt1*Δ *cdc50*Δ mutants exhibited hypersensitivity to the calcineurin inhibitor FK506, highlighting the critical role of flippase in calcium regulation (Figure 7B). The equivalent sensitivity observed in *apt1*Δ mutants further suggests that Apt1 and Cdc50 function together to regulate calcium/calcineurin signalling (Figure 7B).

**Figure 7.**
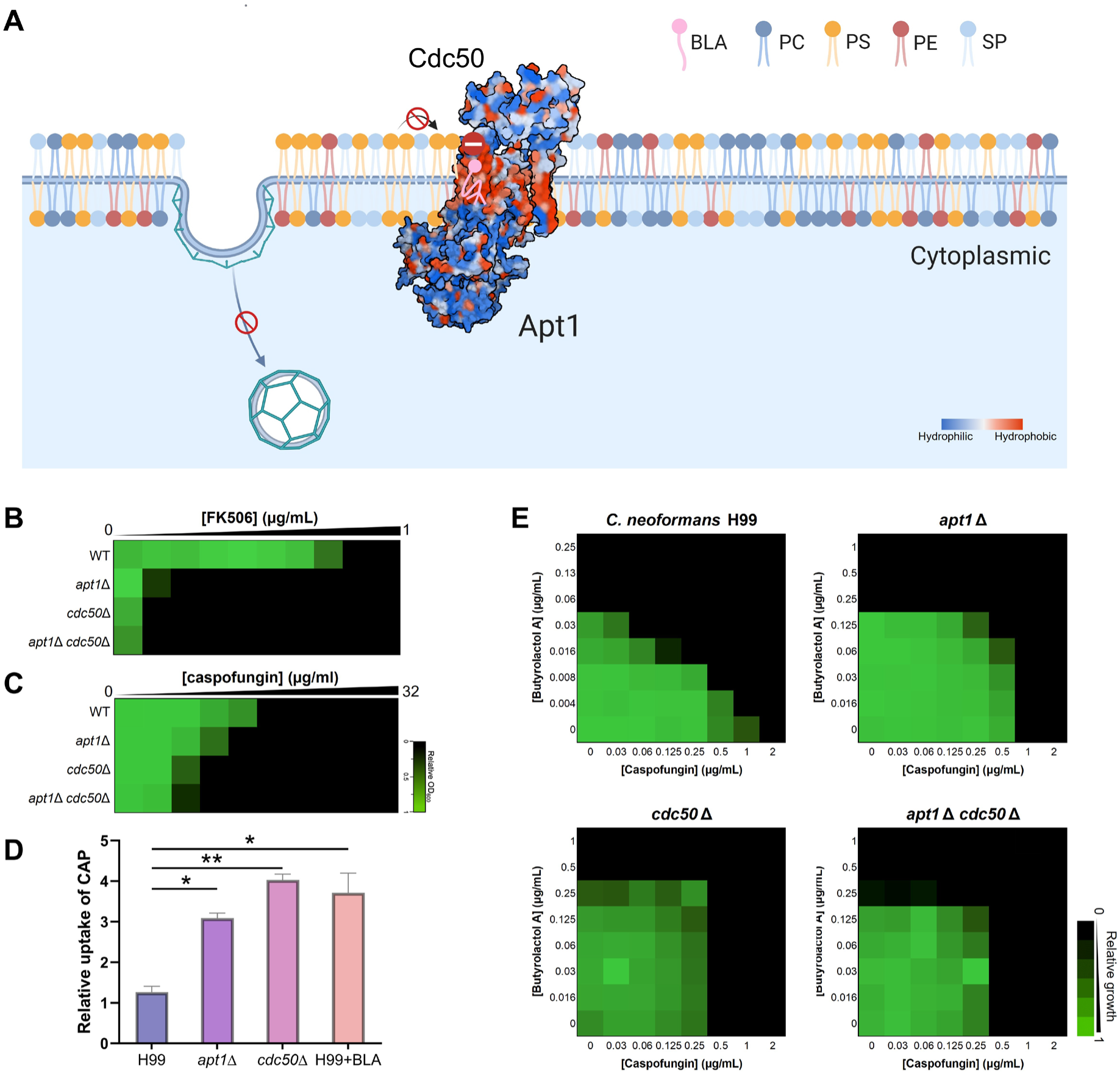
Butyrolactol A (BLA) potentiates the fungicidal activity of caspofungin against Cryptococci by inhibiting the Apt1–Cdc50 complex. **(A)** Schematic illustrating the proposed mechanism of action for BLA. The misloading of BLA blocks the entrance of the phospholipid substrate, resulting in impaired membrane asymmetry, which disrupts membrane organization and vesicle budding, ultimately blocking secretory and endocytosis pathways. Color coding is indicated within the figure. **(B)** Susceptibility of FK506 against *C. neoformans* H99 and flippase-deficient (*cdc50*Δ, *apt1*Δ, and *apt1*Δ *cdc50*Δ). A broth microdilution assay was performed in RPMI 1640 medium, and growth was measured by absorbance at 530 nm after 48 hours at 30 °C. Results are analyzed and depicted as described in Figure 5F. **(C)** Susceptibility of caspofungin against *C. neoformans* H99 wild type, *apt1*Δ, *cdc50*Δ and *apt1*Δ *cdc50*Δ mutants. Dose-response assays were performed, and data were analyzed as described in Figure 5F. Relative growth is represented quantitatively by color (see bar for scale). **(D)** Relative caspofungin accumulation in *C. neoformans* H99 wild type, with or without BLA, and in *apt1*Δ and *cdc50*Δ mutants after 15 minutes of treatment. All experiments were performed in biological triplicates, and data are presented as mean ± SEM. Significance was determined using multiple unpaired t-tests, comparing the wild-type without BLA to each group; p-value: *< 0.05, ** < 0.01. **(E)** The synergistic effect of caspofungin and BLA was assessed against *C. neoformans* H99 wild type, *apt1*Δ, *cdc50*Δ, and *apt1*Δ *cdc50*Δ mutants. Checkerboard assays were performed in SDB medium at 30 °C, and growth was measured by absorbance at 530 nm after 48 hours. Biological duplicates were averaged, and measurements were normalized to no-drug controls and depicted as heatmaps. Relative growth is represented quantitatively by color (see scale bar at the bottom right).

Moreover, deletion of *CDC50* and *APT1* resulted in increased susceptibility to caspofungin, along with significantly elevated caspofungin internalization, demonstrating that both Cdc50 and Apt1 influence caspofungin uptake and potency in *C. neoformans* (Figures 7C-7D). However, the effect of *CDC50* was more pronounced than that of *APT1*, suggesting a broader role for *CDC50* beyond acting as a subunit of *APT1*. Similarly, BLA treatment overcame the limitation of caspofungin uptake, indicating that it potentiates caspofungin activity by inhibiting the Apt1– Cdc50 complex (Figure 7D). This mechanism was further evaluated by examining the interaction in deletion mutants, where the synergistic effect was completely lost in *apt1*Δ, *cdc50*Δ, and *apt1*Δ *cdc50*Δ mutants, confirming that BLA’s adjuvant mechanism involves the inhibition of the Apt1– Cdc50 complex (Figure 7E).

## Discussion

The development of antifungal therapies remains a major challenge due to the conserved eukaryotic cellular machinery and biochemistry shared between fungi and their hosts. Echinocandins represent a relatively non-toxic class targeting fungal cell wall synthesis, but their utility is compromised by intrinsic and acquired resistance. To overcome this limitation, we developed a cell-based screening strategy and identified butyrolactol A (BLA), a previously understudied compound, as a potent adjuvant that restores echinocandin efficacy against intrinsically resistant *Cryptococcus neoformans* and multidrug-resistant *Candida auris*.

BLA’s distinct mechanism of action highlights P4-ATPases as promising antifungal targets, offering a strategy to overcome fungal drug resistance and expand the limited antifungal arsenal. Flippases mediate phospholipid transport between bilayer leaflets, which is crucial for membrane asymmetry^77^ and trafficking^78^, vesicle budding,^79^ polarity maintenance,^80^ and cytoskeletal dynamics.^81^ Among the four flippases in *C. neoformans*, Apt1 plays a broader role in survival and virulence,^43, 44, 67, 82^ exhibiting features of different subfamilies of P4-ATPases, Drs2 and Dnf1.^45, 54^ Although Apt1 is evolutionarily closer to Dnf1 (Figure S5C), it functionally resembles Drs2,^45, 54^ possessing conserved residues for PS specificity, including Q111 in the QQ motif for PS recognition and N477 for PS headgroup stabilization (Figure S11).^49, 57^ The Apt1–Cdc50 complex exhibits broad substrate specificity beyond PS (Figure S5D), permitting BLA misloading but not TAG, indicating that a hydrophilic headgroup is essential for transport despite tolerance to structural variations, enlightening the optimization of structural derivatives for enhanced potency.

BLA’s killing mechanism requires functional Apt1–Cdc50 to form a toxic complex, disrupting membrane asymmetry, architecture, vesicle trafficking, and cytoskeletal organization, ultimately causing cell lysis and death. This mechanism parallels well-known cases where active protein-ligand complexes lead to drug action, while loss-of-function mutations in target proteins confer resistance. For instance, the topoisomerase I (Top1) inhibitor camptothecin induces cytotoxicity by trapping Top1 in enzyme-DNA cleavage complexes, thereby interfering with DNA replication and transcription.^66^ Similarly, calcineurin, a Ca^2+^/calmodulin-dependent phosphatase, is targeted by immunophilin-immunosuppressant complexes, such as cyclophilin A-cyclosporin A (CyPA-CsA) and FKBP12-FK506. Mutations in CyPA or FKBP12 confer resistance to CsA and FK506, respectively.^83, 84^ In addition, rapamycin also targets FKBP12 but forms distinct protein-ligand complexes that disrupt TOR signaling pathways, leading to immunosuppression through a distinct mechanism.^85, 86^ These examples underscore the importance of protein-ligand complexes as the active entities, rather than the small ligands alone.

Our data reveal an unexpected layer of selectivity in BLA’s mechanism of action: beyond direct inhibition of its protein target, BLA toxicity depends on a defined membrane lipid environment shaped by glucosylceramide (GlcCer). The identification of a *CNAG_03870* nonsense mutation, conferring extreme resistance only in the absence of Apt1, highlights a synthetic route to resistance wherein dual disruption of flippase function and GlcCer biosynthesis abolishes susceptibility. This dual requirement suggests that BLA traps the Apt1–Cdc50 complex within a GlcCer-enriched membrane domain, and that the membrane lipid context is not merely permissive but essential for drug activity. These findings support a broader paradigm in which the efficacy of membrane-targeting agents is governed by both protein conformation and local lipid architecture. The dual requirement of BLA for both a functional flippase complex and a defined membrane lipid context has notable therapeutic implications. This mechanism confers high specificity for fungal targets, reducing the likelihood of off-target toxicity in mammalian cells, where the composition and regulation of GlcCer and P4-ATPases differ significantly.

Although Apt1 and Cdc50 form a stable complex that supports flippase activity and display overlapping phenotypes upon gene deletion, a significant growth defect was observed only in *cdc50*Δ mutants.^44, 45^ Nonetheless, spontaneous mutations conferring BLA resistance were found exclusively in *apt1* across independent mutants, suggesting that *cdc50* mutations likely incur a substantial fitness cost. As the sole CDC50 family member in *Cryptococcus*, Cdc50 may have broader roles beyond chaperoning, including the regulation of salt stress, acidic pH, iron metabolism, membrane integrity, and cytoplasmic calcium homeostasis.^23, 44^ The role of Cdc50 in the physiology and pathogenesis of *C. neoformans* warrants further investigation. Recent research indicates that Cdc50 is essential for cryptococcal resistance, although the underlying mechanisms remain unclear.^22, 87^ A synthetic peptide designed to block Cdc50 has shown promise in sensitizing *Cryptococcus* to caspofungin, despite a relatively high FICI, highlighting the potential for rational drug design targeting this subunit.^24^ Our high-resolution structure of the Apt1–Cdc50–BLA complex offers a foundation for designing dual-subunit inhibitors that mimic native lipid substrates and selectively disrupt fungal flippase function.

### Limitations of the Study

This study has several limitations. First, methanol was utilized for extraction due to its analytical compatibility and broad polarity coverage, but highly hydrophobic compounds may have been missed. Second, potentiator screening was performed at a single concentration, which may have overlooked dose-dependent effects or weaker hits. Third, while the *Candida auris* skin colonization model offers a relevant mammalian context, it does not reflect the pathophysiology of disseminated *Cryptococcus neoformans* infection; systemic models will be needed to evaluate efficacy in that setting. Fourth, although BLA showed low cytotoxicity in human cell lines, including THP-1 immune cells, its impact on flippase-dependent membrane dynamics and immune signaling remains untested. Further studies using membrane asymmetry assays, trafficking models, and immunological readouts are needed to assess potential off-target effects, particularly on the ATP8B1–CDC50A complex. In vitro binding assays with purified human flippases will also be necessary to confirm the lack of inhibition observed in cells. While most mechanistic studies— including ours—use detergent-solubilized proteins, reconstitution in liposomes could improve assay fidelity. Fifth, analysis of sterol biosynthesis was limited to *Candida albicans*. Given species-specific differences in sterol regulation and membrane architecture, studies in *C. neoformans* ERG mutants will help clarify how BLA interacts with sterol homeostasis and flippase activity. Finally, the pharmacokinetic properties of BLA remain to be determined and will be essential for translational development.

## RESOURCE AVAILABILITY

### Lead contact

Further information and requests for resources should be directed to and will be fulfilled upon reasonable request by the lead contact, Gerard D. Wright.

### Materials availability

Strains and plasmids generated in this study are available upon request to the lead contact.

### Data and code availability

The Whole Genome Shotgun project of *Streptomyces* sp. WAC8370 has been deposited at DDBJ/ENA/GenBank under the accession JBIEOC000000000. The version described in this paper, JBIEOC010000000 (BioProject accession PRJNA1166230), is included in the key resources table and is publicly available as of the publication date.

The NMR data for butyrolactol A have been deposited in the Natural Products Magnetic Resonance Database (NP-MRD; www.np-mrd.org) and can be found at NP0004856 (https://np-mrd.org/natural_products/NP0004856).

The high-resolution electrospray ionization mass spectrometry data (HR-ESI-MS and MS/MS) for butyrolactol A have been deposited in the Global Natural Products Social Molecular Networking (GNPS) database and are publicly available upon publication [doi:10.25345/C5RR1Q07C]. The dataset is licensed under CC0 1.0 Universal (CC0 1.0). **Pre-publication access (FTP):** ftp://MSV000098469@massive-ftp.ucsd.edu. **Permanent access (upon publication)**: ftp://massive-ftp.ucsd.edu/v10/MSV000098469/.

The cryo-EM 3D maps of the *C. neoformans* E2P state of Apt1–Cdc50 with bound butyrolactol A and the E1 state of Apt1–Cdc50 have been deposited in the Electron Microscopy Data Bank under accession codes EMD-70618 and EMD-47339, respectively. The atomic models of the E2P state of Apt1–Cdc50 with bound butyrolactol A and the E1 state of Apt1–Cdc50 have been deposited in the Protein Data Bank under accession codes 9OMV and 9DZV, respectively. The EM data are available from the lead contact upon reasonable request.

Any additional information required to reanalyze the data reported in this paper is available from the lead contact upon request.

## ACKNOWLEDGMENTS

This work was supported by the Canadian Institutes for Health Research [foundation grant FDN148463 and project grant PJT190298 to G.D.W.] and the U.S. National Institutes of Health grant R01CA231466 (to H.L.) and the Van Andel Institute (to H.L.).

Studies in the Heitman lab (MH, ZB, YC, AFA, SS, and JH) were supported by NIH/NIAID R01 grants AI39115-27, AI050113-20, AI170543-02, AI172451-02, and AI133654-07, and NIH predoctoral fellowship F31 AI150120. J.K. and G.W. are fellows and L.C. and J.H. are fellows and co-directors for the CIFAR program Fungal Kingdom: Threats & Opportunities. J.K. is supported by NIH/NIAID R01 grant AI053721-20.

L.E.C. is supported by the Canadian Institutes of Health Research (CIHR) Foundation grant (FDN-154288) and a National Institutes of Health (NIH) R01 grant (R01AI127375). L.E.C. is a Canada Research Chair (Tier 1) in Microbial Genomics & Infectious Disease and co-Director of the CIFAR Fungal Kingdom: Threats & Opportunities program. CIFAR Catalyst Grants on BLA: CF-0185, CF-0116; Cowen, L. (University of Toronto); Heitman, J. (Duke University); Wright, G. (McMaster University); and Boone, C (University of Toronto).

This research was supported by a Tier 1 Canada Research Chair award and a Foundation grant from the Canadian Institutes of Health Research (CIHR; FRN 143215) to E.D.B.; CIHR funding MOP-119391 to R.T.; and a CIHR grant PJT-156067 to L.M.

The authors gratefully acknowledge the Centre for Microbial Chemical Biology (CMCB) for instrumentation and support, with special thanks to Tracey Campbell, Susan McCusker, and Nicola Henriquez. We also thank the Canadian Centre for Electron Microscopy (CCEM), in particular Marcia Reid, and the McMaster Centre for Advanced Light Microscopy (CALM), with appreciation to Mouhanad Babi and Joao Pedro Bronze de Firmino for their technical assistance. We are grateful to the Nuclear Magnetic Resonance (NMR) Facility at McMaster, especially Bob Berno, for his help in characterizing the stereochemistry of BLA. We also acknowledge the support of the John Mayberry Histology Facility, with special thanks to Mary Jo Smith and Xiaoxing Ma. Cryo-EM data were collected in the David Van Andel Advanced Cryo-Electron Microscopy Suite at the Van Andel Institute. We thank Dr. Gongpu Zhao and Dr. Xing Meng for their facilitation of data collection. We are deeply grateful to Dr. Seok-Yong Lee, Dr. Jiyong Hong, Dr. Kenichi Yokoyama, Dr. Marco Dias Coelho, Dr. Vikas Yadav, Dr. Jun Huang, and Ziyan Xu for their invaluable guidance, expertise, and support during X.C.’s time in the Heitman Lab at Duke University. We thank Dr. Kevin Kavanagh and Dr. Zhongle Liu for their valuable advice and discussion on the histopathological analysis of *Galleria mellonella*, Dr. Dirk Hackenberger and Maya George for their assistance with bioinformatic analyses, and Victoria Lange for technical support with immune cell culture. Finally, we are grateful to Linda Ejim, Dr. David Sychantha, Carlos Barba Bazan, Dr. Nicholas Waglechner, Dr. Grace Yim, Dr. Shawn French, Dr. Elizabeth Culp, Dr. Matt Surette, and Dr. Haley Zubyk for their insightful discussions and ongoing support.

## AUTHOR CONTRIBUTIONS

X.C. and G.D.W. led the project, designed experiments, interpreted data, and wrote the manuscript, with input from all authors. X.C., M.S. and G.D.W. conceived the study. H.D.D. and H.L. designed the cryo-EM study, with X.C. purifying the compound and generating the Apt1–Cdc50 overexpression construct. H.D.D. purified the protein complex, conducted cryo-EM, built atomic models, constructed the expression constructs for ATPase activity assays and measured ATPase activity. M.H. and X.C. designed and conducted the BLA-resistant mutant generation study. M.H. and S.S. performed the genetic cross-assay and analyzed the data, with S.S. generating the double deletion mutants *apt1Δ apt2Δ* and *apt2Δ cdc50Δ*. X.C. prepared the genomic DNA, and A.G. performed genome sequencing and conducted data analysis. M.S. designed, conducted and analyzed the screen, and M.A.C. and X.C. performed additional statistical analysis. K.K., W.W., M.S. and X.C. performed BLA structural elucidation. E.P., B.Y., and N.R. performed dose-response assays on *C. albicans* mutants. S.C. and X.C. conducted the *C. elegans* animal study. G.H. constructed the DsRed-Apt1p. X.C. and Z.B. generated single-point mutations in cryptococcal *APT1*. A.F., Y.C., and A.F.A. performed the murine infection studies. R.T., X.C. and A.B.Y.G. performed wide-field microscopy. X.C. designed and performed all other experiments. R.T., L.T.M., E.D.B., J.W.K., B.K.C., L.E.C., J.H., H.L., and G.D.W. provided resources.

## DECLARATION OF INTERESTS

L.E.C. is a co-founder and shareholder in Bright Angel Therapeutics, a platform company for the development of novel antifungal therapeutics. E.D.B is the CEO and L.E.C. and G.D.W. are Science Advisors for Kapoose Creek Bio, a company that harnesses the therapeutic potential of fungi. All other authors have no competing interests to declare.

## SUPPLEMENTAL INFORMATION

Document S1. Figures S1–S11, Tables S1–S10, Notes S1, S2 and supplemental references.

## Notes

### Summary of Updates

The section entitled Butyrolactol A synergizes with caspofungin in Cryptococcus spp. and resistant Candida auris has been updated to include a detailed analysis of fungal burden in the Galleria model, incorporating both histological studies and quantitative CFU data. In addition, we have introduced a novel murine model of Candida auris infection, further supporting the in vitro efficacy findings and our cytotoxicity assessments, which demonstrated no significant effects on mammalian cells. Figures 2, 4, 5, and 7 have been revised accordingly, and the supplemental files have been updated.

