## Supplementary material for "Butyrolactol A potentiates caspofungin efficacy against resistant fungi via phospholipid flippase inhibition": Figures S1-S11, Table S1-S10, Note S1-S2

Supplementary Figure 1

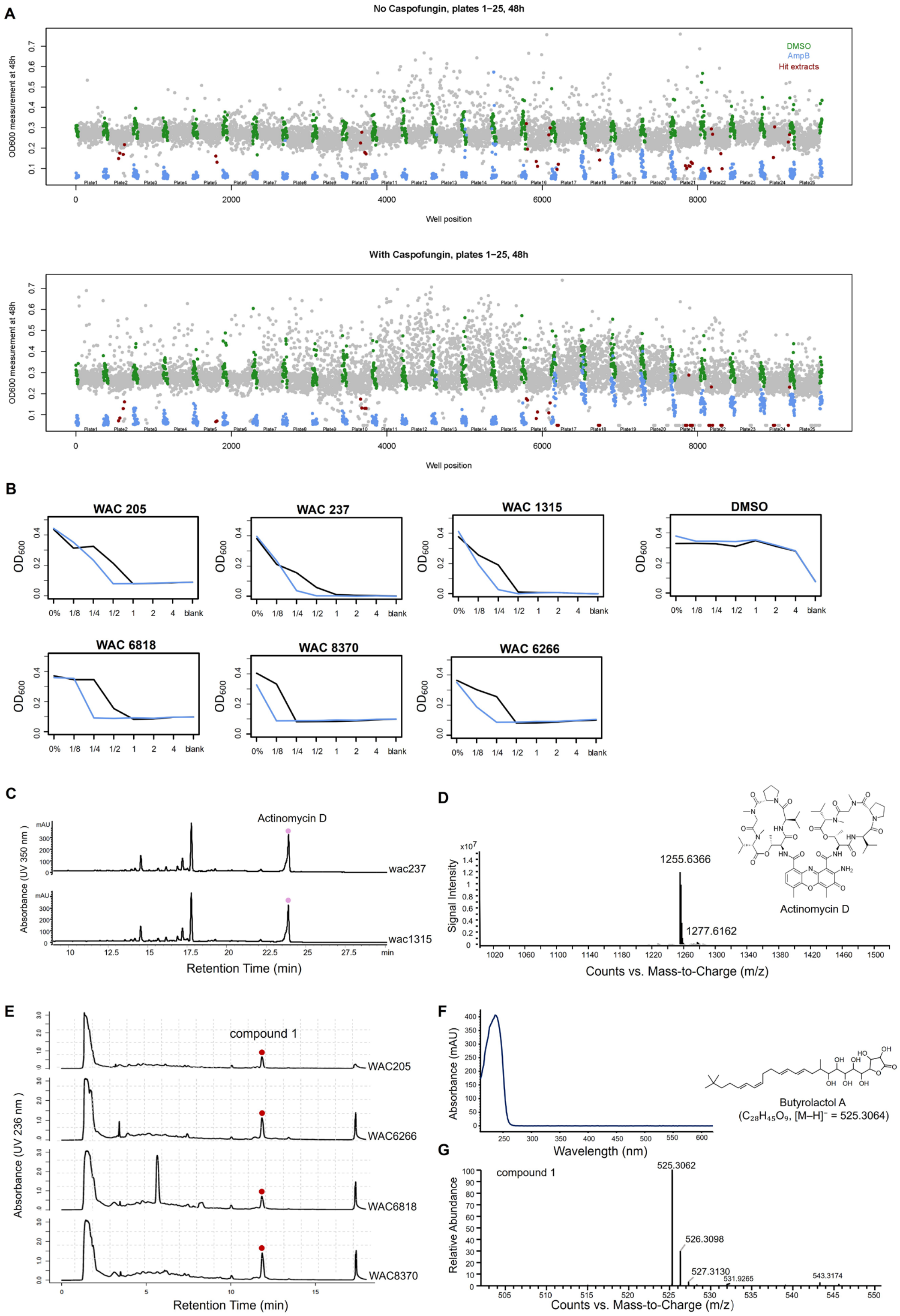


**Figure S1: A high-throughput screen identifies potentiators of caspofungin against *C. neoformans.* Related to Figure 1A.**

(A) Scatter plots displaying the screening results for caspofungin potentiators against *C. neoformans* H99. Growth inhibition caused by compounds in the absence (upper panel) and presence (lower panel) of caspofungin at 1/8th of the minimal inhibitory concentration (MIC) is shown by the $OD_{600}$ measurements (*y*-axis) of *C. neoformans* H99 after treatment with each crude extract from the natural product library (*x*-axis). Red circles highlight compounds classified as active, which inhibited cryptococcal growth by at least 80% in the presence of caspofungin compared to the effect of the compound alone (defined as 2 median absolute deviations below the diagonal). Amphotericin B (AmpB) was used as an antifungal control as was present on each assay plate (blue dots). (B) Six crude extracts from WAC strains were confirmed to potentiate caspofungin activity against *C. neoformans* H99 by comparing dose-response curves with (blue line) and without (black line) caspofungin (5 µg/mL). Growth was measured using absorbance at 600 nm after 48 hours of incubation in RPMI-1640 medium at 30°C. The *x*-axis represents the relative concentration of crude extracts, and the *y*-axis shows pathogen growth. The DMSO control (upper right) did not affect growth in the presence of a sub-inhibitory concentration of caspofungin. (C) HPLC profile of the metabolic products from two actinomycin D-producing strains WAC237 and WAC1315. Bioactivity-guided purification indicated the activity was associated with the compound marked by pink dots, later identified as actinomycin D. (D) Mass spectrum and structure of the purified actinomycin D. (E) HPLC profile of the metabolic products from four hit-producing strains WAC0205, WAC6266, WAC6818, and WAC8370, with compound **1** linked to bioactivity. The red dot indicates the UV absorbance peak corresponding to compound **1**. (F) UV absorbance spectrum of butyrolactol A. UV spectrum of purified butyrolactol A recorded in acetonitrile, showing a maximum absorption at 236 nm and a shoulder at 228 nm. The spectrum was acquired using a diode-array detector (DAD) following HPLC separation (retention time = 14.13 min). (G) Bioactivity-guided purification and identification of butyrolactol A (BLA). The structure (upper) and mass spectrum (lower) of BLA are shown.

Supplementary Figure 2

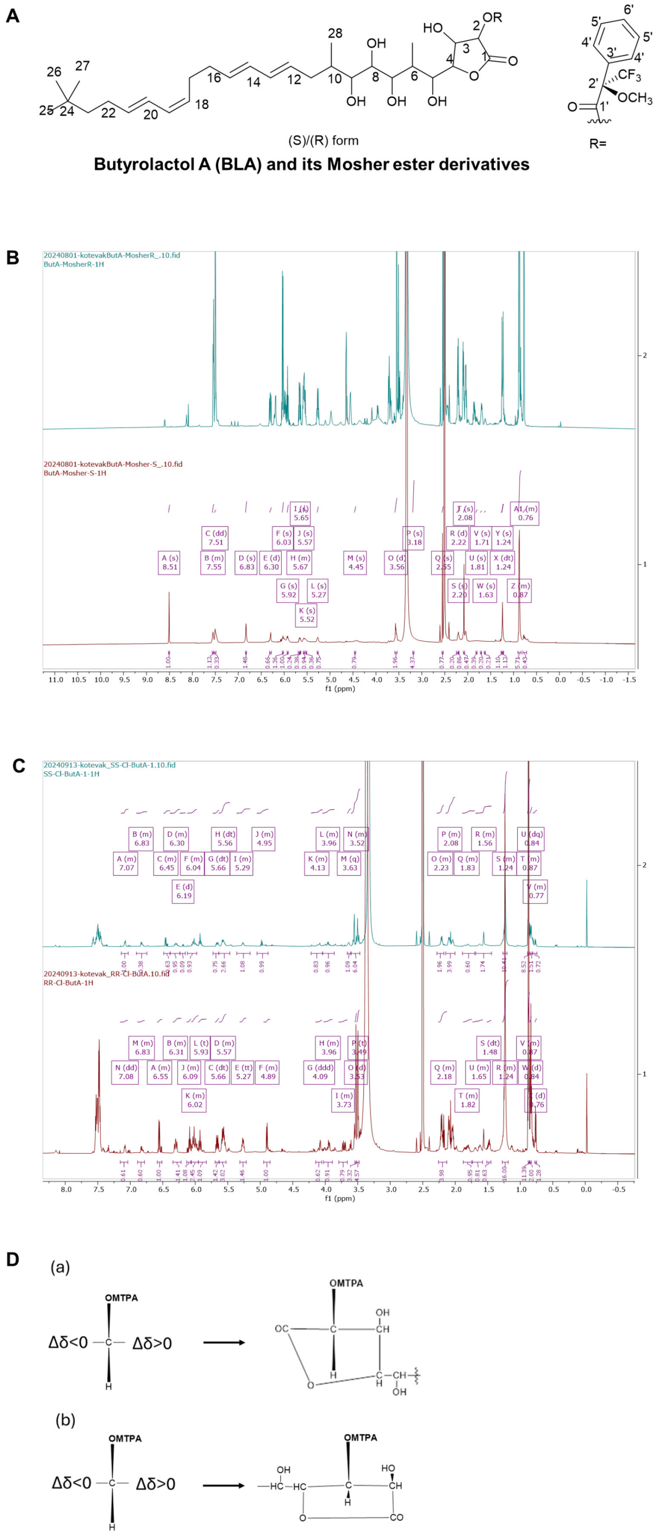


**Figure S2: [1]H-NMR spectra of modified BLA. Related to Figure 1C.**

(A) Structural representation of butyrolactol A (BLA)–MTPA esters used for Mosher analysis. Carbon positions correspond to the assignments listed in Table S1-S3. (B) $^{1}$H NMR overlay of butyrolactol A (BLA) derivatives reacted with R-(−)-MTPA-Cl (top) and S-(+)-MTPA-Cl (bottom) in DMSO-$d_6$, used for Mosher ester analysis. (C) $^{1}$H NMR overlay of butyrolactol A (BLA) diester derivatives prepared by reaction with R-(−)-MTPA-Cl (top) and S-(+)-MTPA-Cl (bottom) in DMSO-$d_6$. These spectra were used to determine the absolute configuration of BLA by the Mosher ester method. (D) Molecular models of BLA–MTPA monoester and diester derivatives used for absolute configuration assignment. (a) Molecular model of the BLA 2-MTPA monoester showing the spatial orientation of protons used in $\Delta\delta$ analysis. (b) Molecular model of the BLA 2,3-MTPA diester confirming the (R, R) absolute configuration at C-2 and C-3. $\Delta\delta$ values from $^{1}$H NMR shifts between diastereomeric esters support this stereochemical assignment according to the modified Mosher method.

Supplementary Figure 3

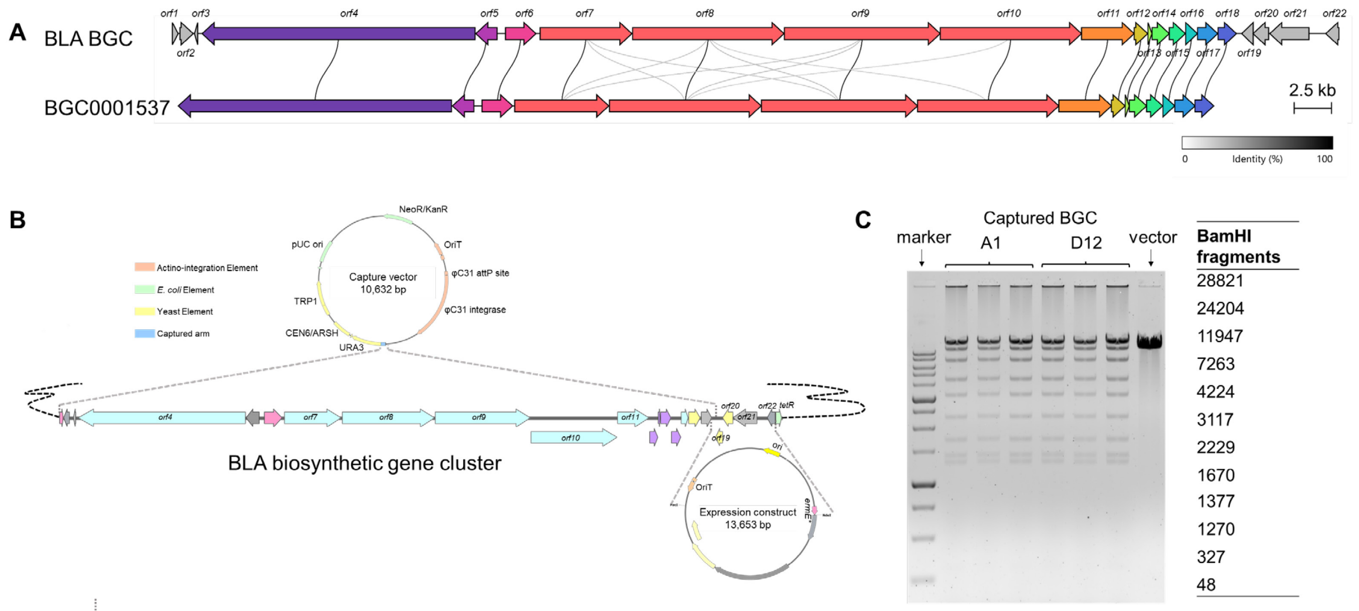


**Figure S3. Construction and validation of the butyrolactol A biosynthetic gene cluster (BGC). Related to Figure 1B–1D. Related to Figure 1B-1D.**

(A) Comparative analysis of the butyrolactol A (BLA) biosynthetic gene cluster (BGC) from *Streptomyces ardesiacus* WAC8370 and a homologous cluster from *Streptomyces* sp. NBRC 110030 (lower; MIBiG accession: BGC0001537). Pairwise alignment was performed using Clinker,[1] revealing conserved gene synteny and extensive homology across core biosynthetic and tailoring genes. Connecting bands indicate homologous regions with ≥30% amino acid identity. Annotated open reading frames (ORFs) in the WAC8370 cluster are listed in Table S4. (B) Strategy for TAR-based capture of the BLA BGC and physical map of the capture construct. A 71,401-bp genomic fragment was recombined into the pCAP03 vector using transformation-associated recombination (TAR), resulting in an 82-kb construct designated pCAP03-BLA BGC. The downstream operon (*orf19–orf22*) was separately cloned into pIJ10257 at *NdeI* and *PacI* sites, excluding the adjacent TetR-family repressor gene to avoid potential transcriptional repression. (C) Restriction digest analysis of pCAP03-BLA BGC using *BamHI*, isolated from two independent yeast transformants (A1 and D12) and propagated in *E. coli* Top10. The expected restriction pattern confirmed successful cloning of the 71-kb genomic region, and the 82-kb construct was stably maintained in *E. coli*.

Supplementary Figure 4

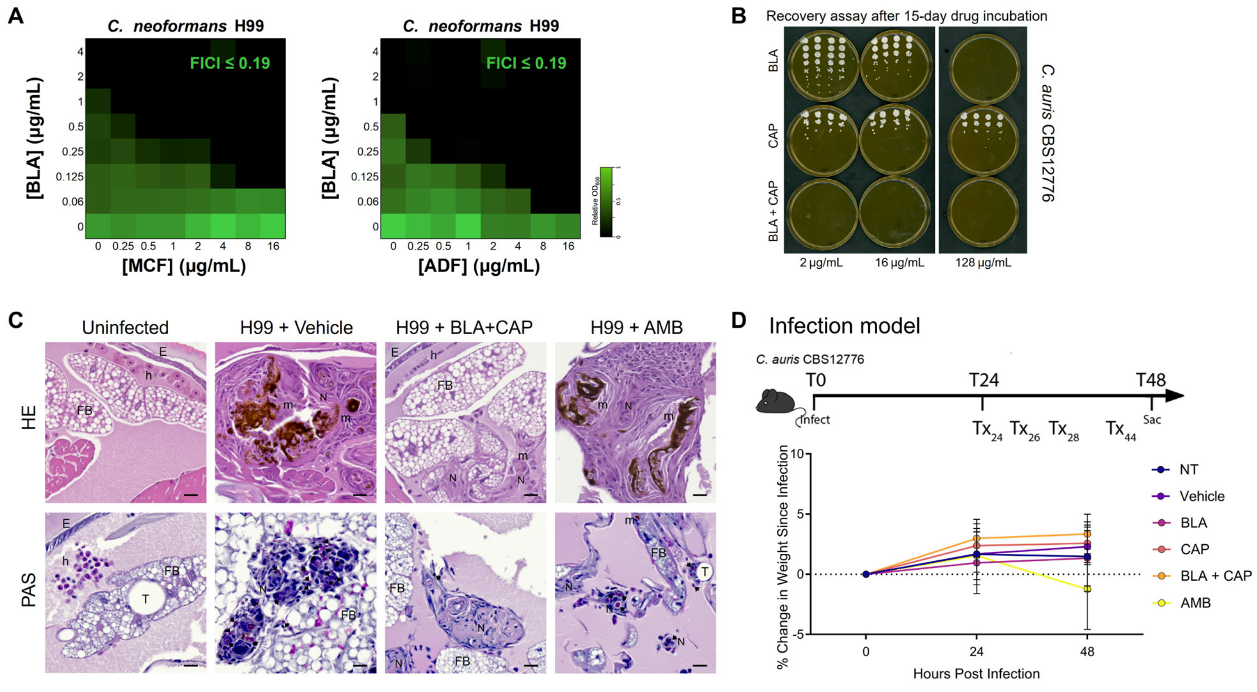


**Figure S4. Butyrolactol A synergizes with echinocandin in *Cryptococcus* spp. and resistant *Candida auris.* Related to Figure 2.**

(A) Checkerboard heatmaps showing growth inhibition of *C. neoformans* H99 by BLA in combination with anidulafungin (ADF) or micafungin (MCF). Cultures were treated in biological duplicates and growth was normalized to no-drug controls. Neither ADF nor MCF exhibited standalone activity at concentrations up to 64 µg/mL. Relative growth is color-coded (see scale bar), and fractional inhibitory concentration index (FICI) values, indicating synergy, are shown in the top-right corner of each matrix. (B) Long-term eradication of *Candida auris* by BLA–CAP combination therapy. *C. auris* CBS12766 cultures were incubated in fresh RPMI 1640 medium containing BLA, caspofungin (CAP), or their combination for 15 days. At the endpoint, cultures were serially diluted ($10^1$–$10^8$), plated on YPD agar, and incubated for 48 h at 30 °C to assess viable colony-forming units (CFUs). BLA monotherapy at 128 µg/mL and the BLA–CAP combination completely eradicated viable fungal cells, with no detectable CFUs at any dilution. Plates without visible colonies were interpreted as below the detection limit and plotted as $\log_{10}$ CFU/mL = 0 for visualization only. Representative plates from four independent biological replicates are shown. Drug concentrations are indicated on the figure. (C) Extended histopathological analysis of *G. mellonella* tissues following *C. neoformans* infection. Representative histological sections of *G. mellonella* larvae 2 days post-infection with *C. neoformans* H99 under indicated treatments. Tissues were stained with hematoxylin and eosin (H&E) or periodic acid–Schiff (PAS) and imaged at 20× magnification. E, epithelial layer; FB, fat body; h, hemocyte; N, hemocyte nodule; m, melanin; T, trachea. Black arrows mark *C. neoformans* yeast cells localized within hemocyte nodules, indicative of immune activation. Scale bar, 20 μm. (D) Body weight monitoring in the *C. auris* murine skin colonization model following topical antifungal treatments. C57BL/6N mice were topically infected with *Candida auris* CBS12766 and treated with vehicle, BLA, caspofungin, BLA + CAP, or amphotericin B (25 mg/mL) at 24, 26, 28, and 44 h post-infection. Body weights were recorded at 0, 24, and 48 h post-treatment initiation. Mice receiving BLA–CAP combination therapy showed no signs of morbidity or weight loss, indicating good tolerability. Data represent mean ± s.e.m.; n = 10 mice per group.

Supplementary Figure 5

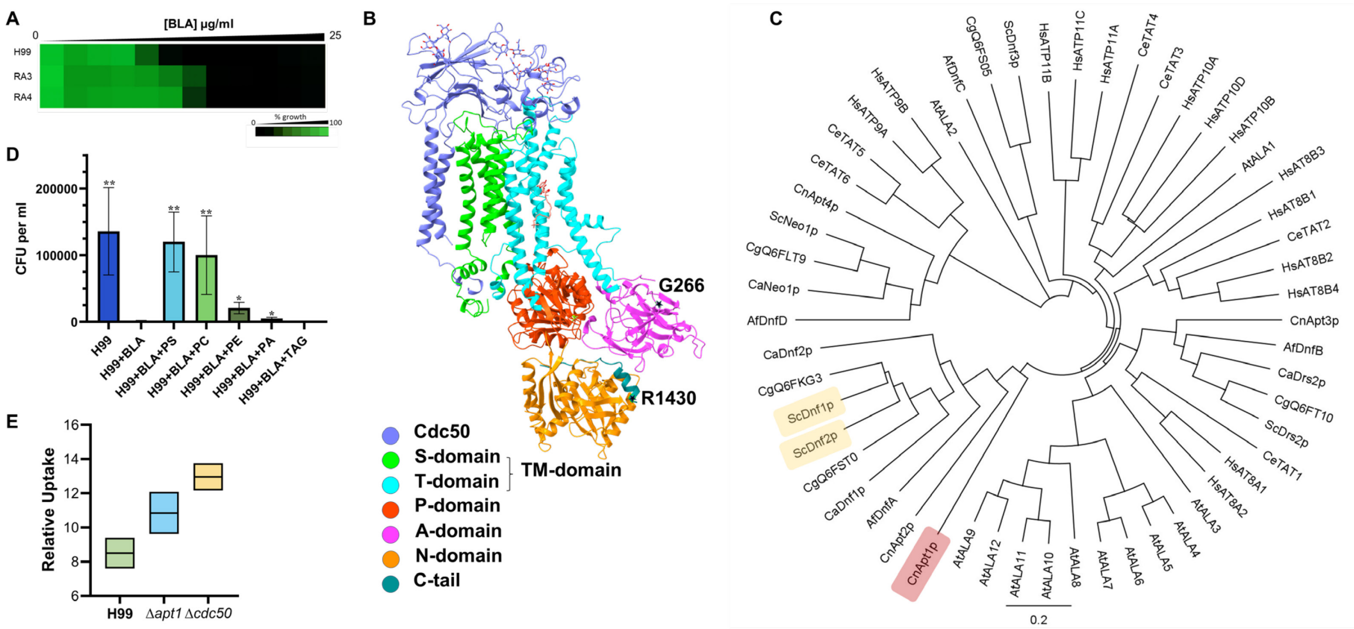

**Figure S5. Identification of a novel mode of action (MOA) of butyrolactol A. Related to Figure 4**

(A) BLA susceptibility test for wild-type *C. neoformans* H99 and lab-generated BLA-resistant mutants R3 and R4. Dose-response assays were performed in RPMI 1640 medium, and growth was measured by absorbance at 530 nm after 48 hours. Optical densities were averaged from duplicate measurements and normalized to no-drug controls. Relative growth is quantitatively represented by color (see scale bar). (B) Mutations found in BLA-resistant mutants RA3 and RA4 are indicated as a black star on the BLA-bound Apt1–Cdc50 E2P structure (PDB ID 9OMV, this study). (C) Phylogenetic tree showing sequences of annotated P4-ATPases from different species, including *C. neoformans* (Cn), *C. albicans* (Ca), *S. cerevisiae* (Sc), *Candida glabrata* (Cg); the *Homo sapiens* (Hs); the model plant *A. thaliana* (At); and the worm *C. elegans* (Ce). The evolutionary history was inferred using the UPGMA method and the Jukes-Cantor genetic distance model. (D) The viability of *C. neoformans* H99 was measured by colony-forming units (CFU) after 24 hours of BLA treatment, with and without supplementation of different lipids: phosphatidylserine (PS), phosphatidylcholine (PC), phosphatidylethanolamine (PE), phosphatidic acid (PA), and triacylglycerol (TAG). Data are presented as mean ± SD of biological triplicates. Statistical significance was determined using multiple unpaired t-tests, comparing each condition with the BLA treatment control. *p-value: * < 0.05, ** < 0.01*. (E) Relative intracellular concentration of amphotericin B accumulated in *C. neoformans* H99 and *Δapt1* and *Δcdc50* mutants were measured after 15 minutes of treatment. Data are presented as mean ± SD of biological triplicates.

### Supplementary Figure 6

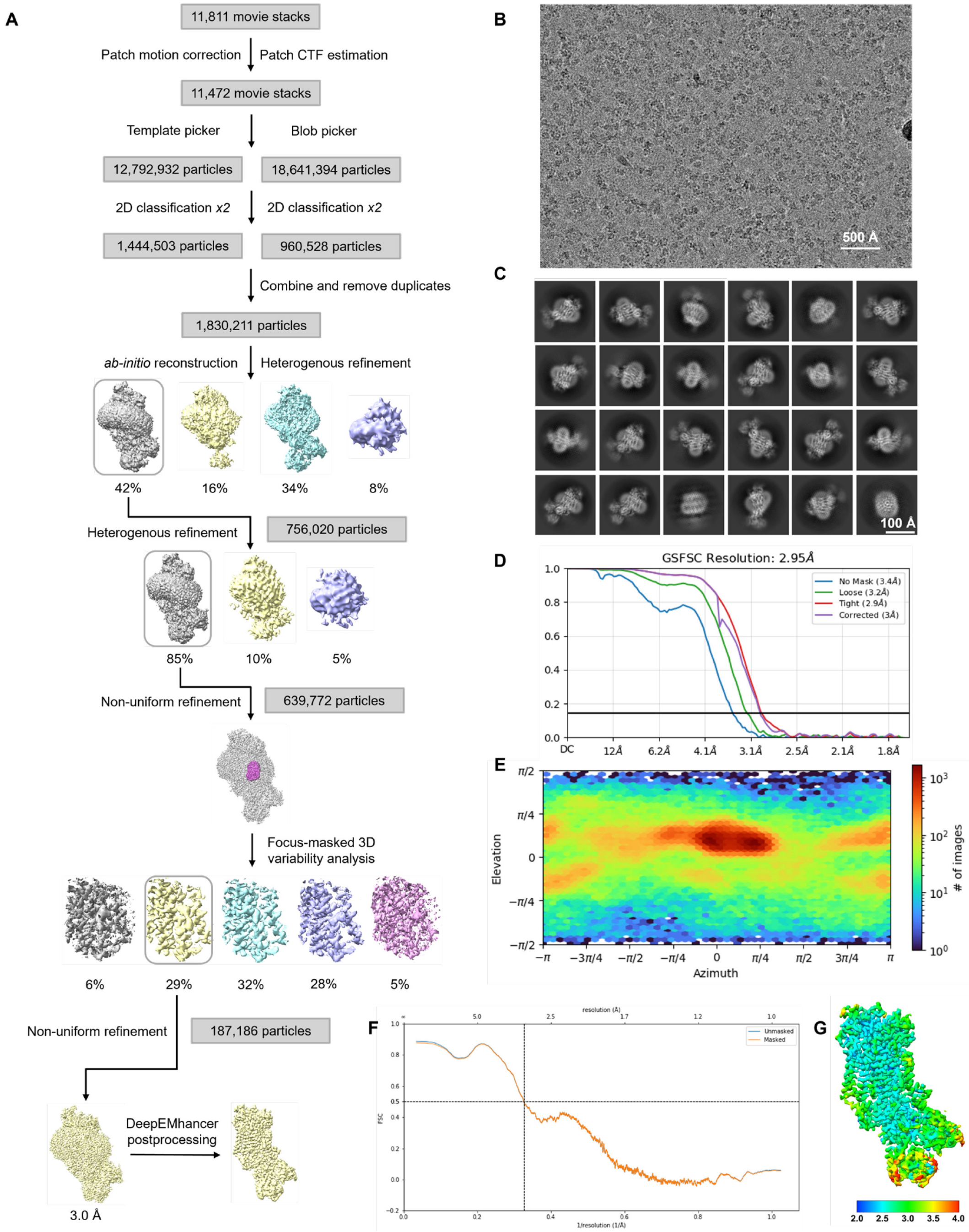


**Figure S6. Cryo-EM structural determination of Apt1–Cdc50 bound with BLA in the E2P state.**

(A) Data processing workflow. (B) Representative raw micrograph. A total of 11,811 such micrographs were recorded. (C) Representative 2D classes. (D) Gold-standard Fourier shell correlation (GSFSC) curve for 3D reconstruction of the consensus map. (E) Angular distribution heat map for 3D reconstruction of the consensus map. (F) Model-to-map FSC curve with the resolution indicated at 0.5 threshold. (G) Local resolution estimation map.

### Supplementary Figure 7

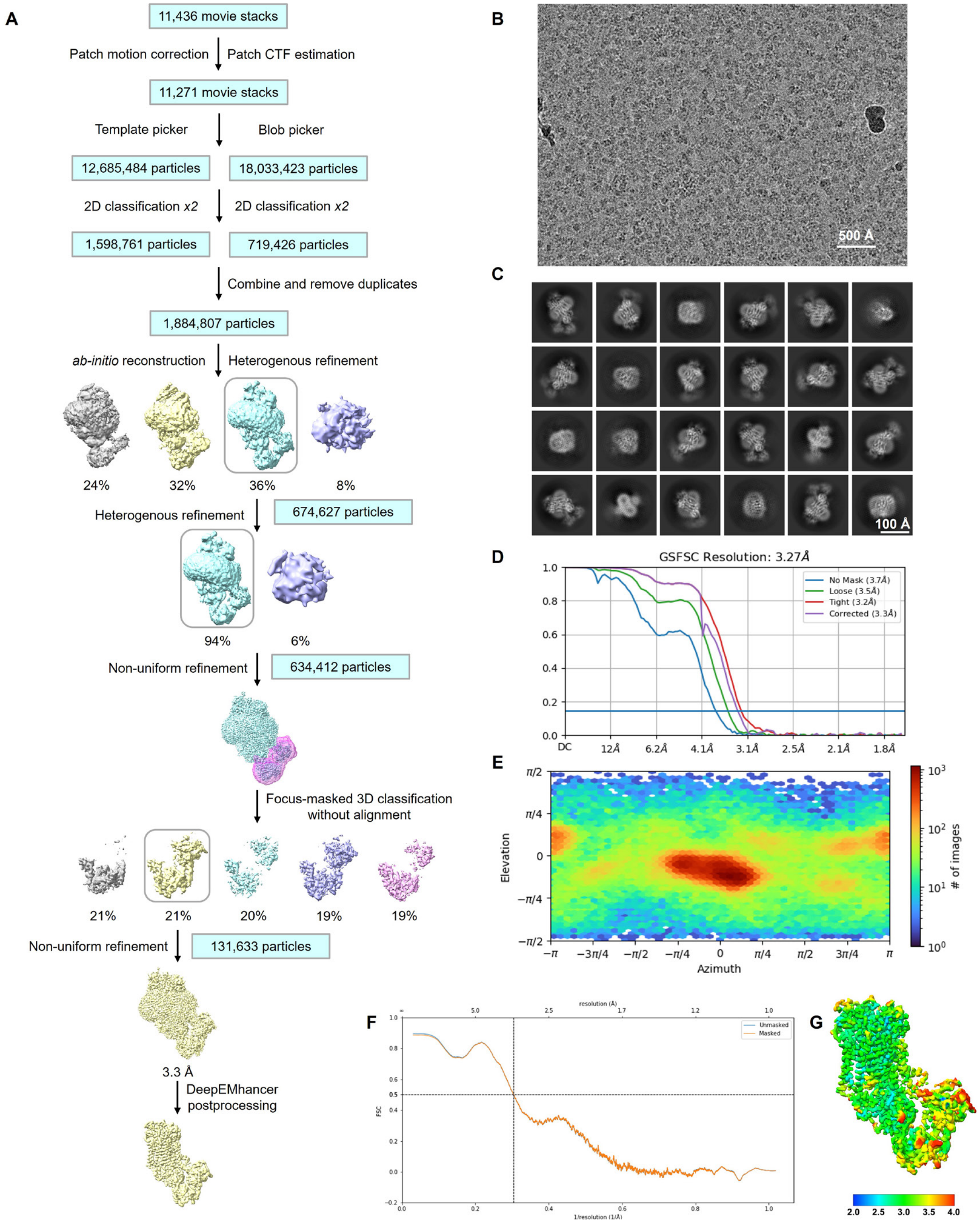


**Figure S7. Cryo-EM structural determination of Apt1–Cdc50 in the E1 state.**

(A) Data processing workflow. (B) Representative raw micrograph. A total of 11,436 such micrographs were recorded. (C) Representative 2D classes. (D) Gold-standard Fourier shell correlation (GSFSC) curve for the 3D reconstruction. (E) Angular distribution heat map for the 3D reconstruction. (F) Model-to-map FSC curve with the resolution indicated at 0.5 threshold. (G) Local resolution estimation map.

**Supplementary Figure 8**

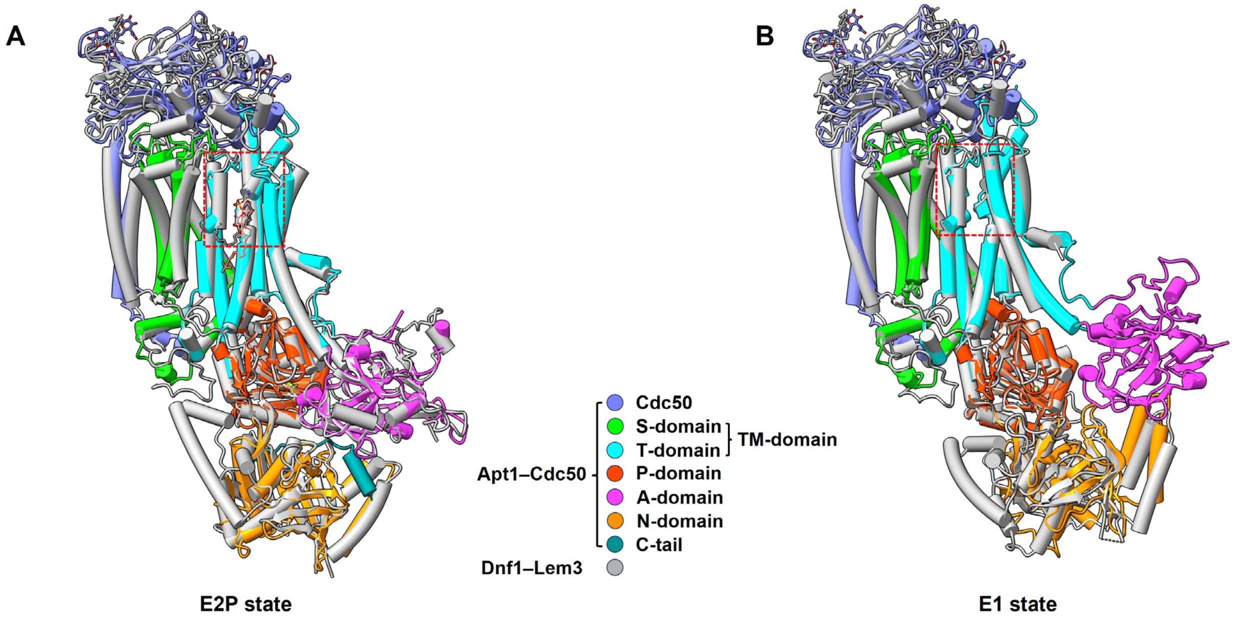


**Figure S8. Superimposition of *C. neoformans* Apt1–Cdc50 and *S. cerevisiae* Dnf1–Lem3. Related to Figure 5B, 5C, 5D and 5E.**

(A) Superimposition of BLA-bound Apt1–Cdc50 in E2P state shown in color (PDB ID 9OMV, this study) and PC-bound Dnf1–Lem3 in E2P state shown in gray (PDB ID 7KYC). The lipid entry site is boxed in red. BLA is shown in salmon sticks and PC is shown in gray sticks. (B) Superimposition of Apt1–Cdc50 in E1 state shown in color (PDB ID 9DZV, this study) and Dnf1–Lem3 in E1 state shown in grey (PDB ID 7KY6). The lipid entry site is boxed in red.

**Supplementary figure 9**

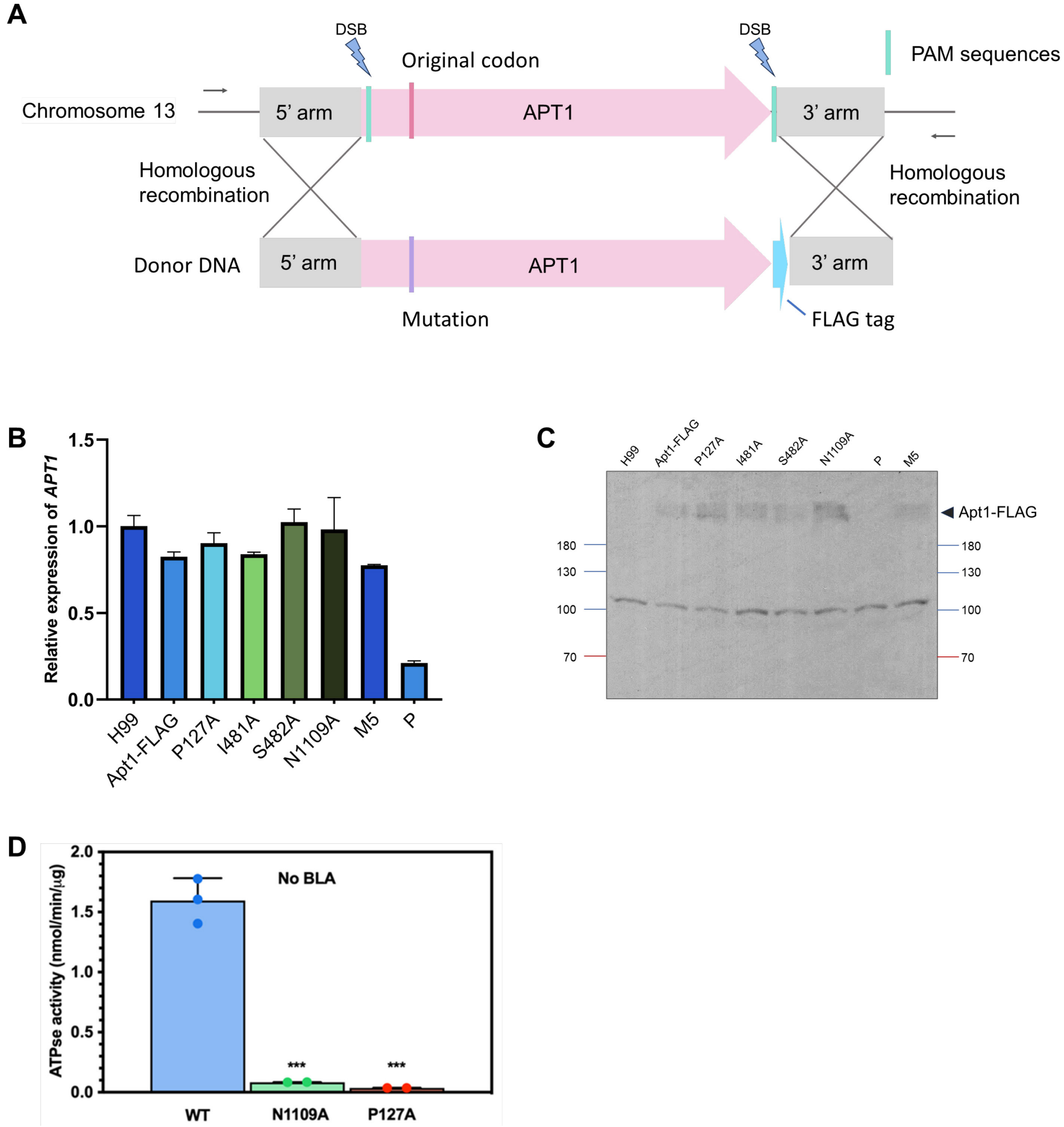


**Figure S9. Mutagenesis study of the BLA binding residues in Apt1. Related to Figure 5F and 5G.**

(A) Schematic representation of the transient CRISPR (clustered regularly interspaced short palindromic repeat)-Cas9 coupled with Electroporation (TRACE) system used for site-directed modification of *APT1* via homologous recombination (HR). The diagram shows the donor DNA and genomic DNA region targeted for *APT1* modification. The donor DNA includes a FLAG-tag at the C-terminus of Apt1, synonymous mutations in the 5' and 3' protospacer-adjacent motif (PAM) and gRNA sequence, as well as either a wild-type codon or an alanine substitution in the designated binding residues. The gRNA target site, indicated by a turquoise box, marks the location of a double-strand break (DSB) induced by Cas9 to facilitate HR via a double cross-over between the donor DNA and the genome. Primers used for PCR verification and sequencing are represented by arrows. (B) qRT-PCR analysis of *APT1* mRNA expression in the *C. neoformans* H99 wild type (WT) and mutant strains. Relative expression levels of *APT1* in WT and mutants were normalized to the housekeeping gene β-actin and presented as fold change relative to WT. Data represent the mean ± standard error (SE) from three independent experiments. Statistical significance was determined by Student's t-test, with $^{**}p < 0.05$ indicating significance. The mutant strains include FLAG-tagged Apt1 (Apt1-FLAG) with various single-point mutations, a quintuple mutation (M5: I130A, I134A, I485A, I493A, I1118A), or a disrupted promoter (P). (C) Western blot analysis of Apt1 protein expression in *C. neoformans* H99 WT and mutant strains. The Apt1-FLAG is indicated by the arrow. The labelling of the strains is consistent with panel B. (D) PS-stimulated ATPase activity of WT, N1109A and P127A Apt1, at 0.05 mM PS and no BLA. Data points represent the mean ± SD in triplicate ($n = 3$) for WT and duplicate ($n = 2$) for mutants. Ordinary one-way ANOVA was performed to test the variance and comparisons with WT Apt1 were made with Dunnett's post hoc analysis. $^{***}p < 0.001$.

### Supplementary Figure 10

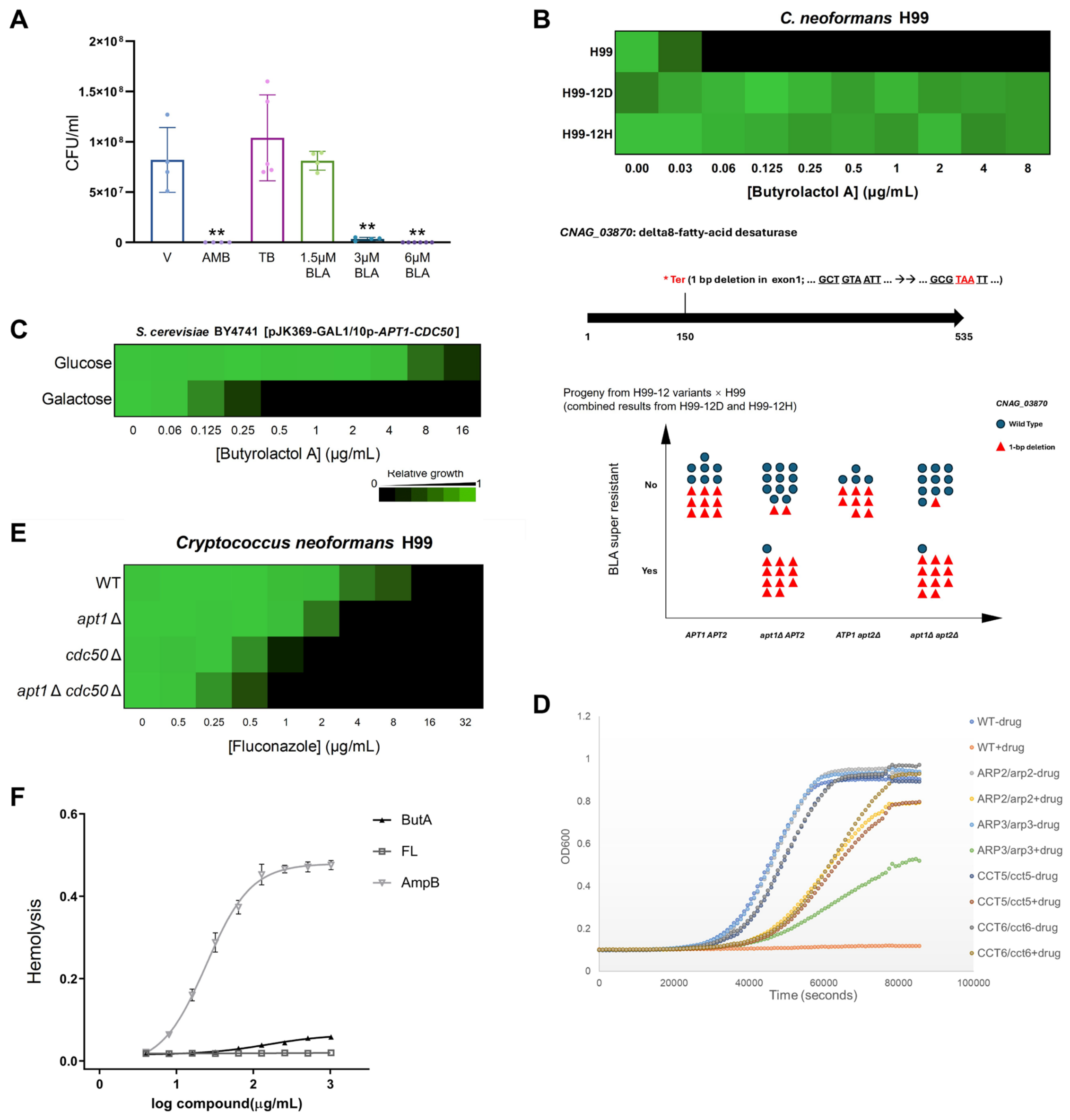

**Figure S10. BLA and antifungal susceptibility of mutants and human cell toxicity. Related to Figure 6.**

(A) **Related to Figure 6A.** Colony-forming units (CFU) of log-phase *C. neoformans* H99 cells were quantified following treatment with Amphotericin B (AMB), terbinafine (TB), butyrolactol A (BLA), or vehicle control (V) in SDB medium. Data are shown as mean ± SD from biological triplicates, with significant differences indicated. Statistical analysis was performed using multiple unpaired t-tests against the vehicle control (***p* < 0.01). (B) A nonsense mutation in *CNAG_03870* confers BLA super-resistance in the absence of *APT1*. Top: Heatmap showing growth of *C. neoformans* H99 and isogenic mutants H99-12D and H99-12H across a range of butyrolactol A concentrations. Strains harboring the *CNAG_03870* 1-bp deletion exhibit marked resistance relative to wild-type H99. Middle: Schematic of *CNAG_03870* (delta8-fatty acid desaturase) showing a 1-bp deletion in exon 1 that introduces a premature stop codon (*Ter). Bottom: Genotype-phenotype association of meiotic progeny from H99-12 variants × H99 crosses. Each dot represents a single progeny, color-coded by *CNAG_03870* genotype (blue circle: wild-type; red triangle: 1-bp deletion). BLA super-resistance was only observed in progeny with both the *apt1*Δ allele and the *CNAG_03870* nonsense mutation, supporting a genetic interaction between *APT1* and glucosylceramide biosynthesis in mediating BLA susceptibility. (C) Two-fold dose-response assays for BLA were performed using *S. cerevisiae* BY4741 *pep4*Δ [pJK369-GAL1/10p-*APT1–CDC50*], under tight regulation of Apt1–Cdc50 expression with galactose or glucose. Growth was measured by absorbance at 600 nm after 48 hours at 30 °C. Data are presented as a heatmap, with colors indicating relative growth normalized to no-drug controls. (D) Dose-response curves were performed for *C. albicans* wild-type and heterozygous actin assembly mutants in the absence and presence of BLA (labelled as 'drug'). Growth was measured by absorbance at 600 nm in YPD medium after 24 hours at 30 °C. Data represent the average of technical triplicates, with each measurement shown as a single dot. Different mutants are represented by distinct colors, with annotations provided on the left of the figure. (E) Susceptibility test of flippase-deficient mutants (*cdc50*Δ, *apt1*Δ, and *apt1*Δ *cdc50*Δ) to fluconazole. Dose-response assays were conducted in RPMI 1640 medium, and growth was measured by absorbance at 530 nm after 48 hours at 30 °C. Optical densities were averaged from duplicate measurements and normalized to wild-type strain growth without compounds. Relative growth is depicted using color, as indicated by the scale bar. (F) Dose-response curve of hemolytic activity of BLA, amphotericin B (AMB), and fluconazole (Flu). Hemolysis was tested using a 0.5% human blood suspension. A 9-point dose-response series was performed with two-fold dilutions. Triton X-100 (1%) was used as a positive control, and DMSO as a negative control. Results are presented as mean ± SD (n = 3).

### Supplementary Figure 11

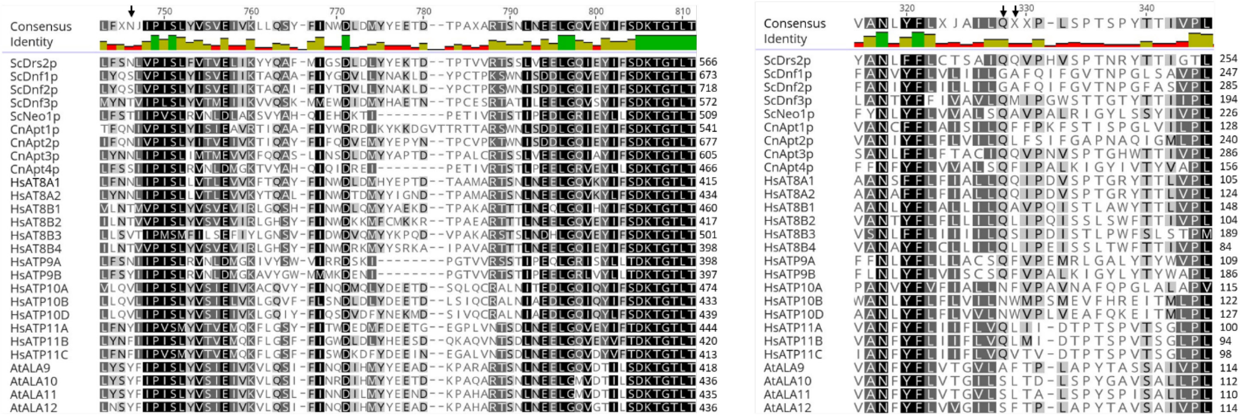

**Figure S11. Aligned fragments of selected P4-ATPases showing residues important for PS selectivity and lipid translocation.** Protein sequences were obtained from Uniprot with the following accession numbers: Yeast/fungi: *Cryptococcus neoformans*: J9VZ19 (Apt1p), J9VQH2 (Apt2p), J9VGP8 (Apt3p), J9VM87 (Apt4p); *Saccharomyces cerevisiae*: P32660 (ScDnf1p), P39524 (ScDrs2p), P40527 (ScNeo1p), Q12675(ScDnf2p), Q12674 (ScDnf3p); *H. sapiens*: Q9Y2Q0 (HsAT8A1), Q9NTI2(HsAT8A2), O43520 (HsAT8B1), P98198 (HsAT8B2), O60423 (HsAT8B3), Q8TF62 (HsAT8B4), O75110 (HsATP9A), O43861 (HsATP9B), O60312 (HsATP10A), O94823 (HsATP10B), Q9P241(HsATP10D), P98196(HsATP11A), Q9Y2G3 (HsATP11B), Q8NB49 (HsATP11C); *Arabidopsis thaliana*: P98204 (AtALA1), P98205 (AtALA2), Q9XIE6 (AtALA3), Q9LNQ4(AtALA4), Q9SGG3 (AtALA5), Q9SLK6 (AtALA6), Q9LVK9 (AtALA7), Q9LK90 (AtALA8), Q9SX33 (AtALA9), Q9LI83 (AtALA10), Q9SAF5 (AtALA11), P57792 (AtALA12). The sequences were aligned with Clustal Omega. Arrowheads above represents the essential residues for PS selectivity and stabilization.

**Supplementary Table 1. $^1$H NMR chemical shift data (ppm, DMSO-$d_6$) for butyrolactol A (BLA) and its Mosher ester derivatives.**

| | BLA | | BLA+ *S*-(+)-MTPA-Cl (R form) | | BLA+ *R*-(−)-MTPA-Cl (S form) | |
|---|---|---|---|---|---|---|
| Carbon # | $\delta_H$ (m, J in Hz) | $\delta_C$ | $\delta_H$ (m, J in Hz) | $\delta_C$ | $\delta_H$ (m, J in Hz) | $\delta_C$ |
| 1 | - | | - | 169.3 | - | 169.3 |
| 2 | 4.25 (dd, J = 9.0, 5.9 Hz, 1H) | 74.03 | **6.08 (m, 1H)** | **69.8** | **6.02 (m, 1H)** | 69.7 |
| 2-OH | 6.04 (d, 1H, 6.4) | - | missing | - | missing | - |
| 3 | 4.16 (td, J = 8.7, 6.1 Hz, 1H) | 72.38 | **4.46 (m, 1H)** | 76.6 | **4.55 (m, 1H)** | 76.5 |
| 3-OH | 5.70 (d, *J* = 6.2 Hz, 1H) | - | **6.11 (m, 2H)** | - | **6.20 (dd, *J* = 16.7, 6.5 Hz, 1H)** | - |
| 4 | 4.32 (dd, *J* = 18.0, 8.2 Hz, 1H) | 79.63 | **4.64 (dd, *J* = 8.4, 1.0 Hz, 1H)** | 80.8 | **4.65 (dd, *J* = 8.4, 1.0 Hz, 1H)** | 80.8 |
| 5 | 3.63 (t, *J* = 8.7 Hz, 1H) | 66.38 | **3.66 (m, 1H)** | **66.4** | **3.67 (m, 1H)** | 66.0 |
| 5-OH | 4.85 (d, *J* = 7.7 Hz, 1H) | - | 4.98 (d, *J* = 7.7 Hz, 1H) | - | 4.98 (d, *J* = 7.7 Hz, 1H) | - |
| 6 | 3.72 (dt, *J* = 10.6, 8.5 Hz, 1H) | 68.47 | 3.72 (m,1H) | 68.4 | 3.72 (m,1H) | 68.3 |
| 6-OH | 4.32 (m,1H) | - | 4.36 (d, *J* = 8.0 Hz, 1H) | - | 4.36 (d, *J* = 7.5 Hz, 1H) | - |
| 7 | 3.72 (dt, *J* = 10.6, 8.5 Hz, 1H) | 68.32 | 3.72 (m,1H) | 68.4 | 3.72 (m,1H) | 68.3 |
| 7-OH | 4.02 (d, *J* = 8.2 Hz, 1H) | - | 4.04 (d, *J* = 8.4 Hz, 1H) | - | 4.05 (d, *J* = 8.4 Hz, 1H) | - |
| 8 | 3.49 (t, *J* = 8.9 Hz, 1H) | 69.14 | 3.50 (m, 1H) | 69.2 | 3.49 (m, 1H) | 69.4 |
| 8-OH | 3.97 (d, *J* = 8.2 Hz, 1H) | - | 3.97 (d, *J* = 8.3 Hz, 1H) | - | 3.97 (d, *J* = 8.3 Hz, 1H) | - |
| 1’ | - | - | | 164.4 | | |
| 2’-OCH3 | - | - | 3.55 (s, 3H) | 55.4 | | |

**Supplementary Table 2. Chemical shift differences ($\Delta\delta = \delta_S - \delta_R$) between diastereomeric BLA-MTPA esters in ppm.**

| Proton | δ BLA-*S*- (+)-MTPA *from R*-(−)-MTPA-Cl (ppm) | δ BLA-*R*-(-)-MTPA from *S*-(+)-MTPA-Cl (ppm) | Δ δ=($\delta_S$-$\delta_R$) (ppm) |
|---|---|---|---|
| 2 (modified) | 6.02 | 6.08 | NA |
| 3 | 4.55 | 4.46 | +0.09 |
| 3-OH | 6.20 | 6.11 | +0.9 |
| 4 | 4.652 | 4.644 | +0.01 |
| 5 | 3.68 | 3.66 | +0.02 |

**Based on the calculated Δδ values, the absolute configuration at C-2 of butyrolactol A is assigned as R.**

**Supplementary Table 3. Chemical shift differences ($\Delta\delta = \delta_S - \delta_R$) for MTPA-(*S)*-BLA and MTPA-(R)-BLA in ppm.**

| Proton # | δ SS-BLA ester (from *R-MTPA-Cl*) (ppm) | δ RR- BLA ester (from *S-MTPA-Cl*) (ppm) | Δ δ= ($\delta_S$-$\delta_R$) (ppm) |
|---|---|---|---|
| 2 | 6.55 | 6.43 | +0.12 |
| 3 | 6.12 | 5.92 | NA |
| 4 | 4.88 | 4.97 | -0.1 |
| 5 | 4.08 | *4.131* | -0.03 |
| 6 | 3.94 | 3.94 | 0 |

**Supplementary Table 4. Annotated putative ORFs in and near the biosynthetic gene cluster of butyrolactol A.**

| orf | Accession no. | size | Gene annotation | Alignment coverage/identity |
|---|---|---|---|---|
| 1 | WP_053639570.1 | 127 | HxlR family HTH-type transcriptional regulator | 100/100 |
| 2 | WP_055469544.1 | 193 | hypothetical protein | 100/96 |
| 3 | WP_030405160.1 | 71 | biotin/lipoyl-binding carrier protein | 100/99 |
| 4 | WP_350892326.1 | 6051 | Type I PKS | 100/99 |
| 5 | WP_030405159 | 473 | Carboxyl transferase domain-containing protein | 100/99 |
| 6 | WP_249956491.1 | 676 | LuxR family HTH-type transcriptional regulator | 100/100 |
| 7 | WP_234353271.1 | 2078 | Type I PKS | 100/98 |
| 8 | WP_055469548.1 | 3359 | Type I PKS | 100/98 |
| 9 | WP_267715824.1 | 3459 | Type I PKS | 100/99 |
| 10 | WP_055469550.1 | 3135 | Type I PKS | 100/99 |
| 11 | WP_053664162.1 | 1169 | Type I PKS | 100/99 |
| 12 | WP_030403676.1 | 301 | 3-hydroxybutyryl-CoA dehydrogenase | 100/99 |
| 13 | WP_030403675.1 | 88 | acyl carrier protein | 100/100 |
| 14 | WP_055469553.1 | 381 | acyl-CoA dehydrogenase | 100/100 |
| 15 | WP_350892282.1 | 356 | HAD-IIIC family phosphatase | 100/100 |
| 16 | WP_030403672.1 | 261 | alpha/beta fold hydrolase | 100/100 |
| 17 | WP_031086102.1 | 448 | MFS-type transporter | 100/100 |
| 18 | WP_108933443.1 | 417 | hypothetical protein | 100/99 |
| 19 | WP_359515283.1 | 253 | ABC transporter permease | 100/99 |
| 20 | WP_078869586.1 | 346 | ABC transporter ATP-binding protein | 100/99 |
| 21 | WP_108933445 | 868 | class I SAM-dependent methyltransferase | 100/100 |
| 22 | WP_053664170.1 | 290 | ketopantoate reductase | 100/99 |

**Supplementary Table 5.** ***In vitro*** **bioactivity assessment of butyrolactol A.**

| **Organism** | **MIC (μg/mL)** | | | | |
|---|---|---|---|---|---|
| | Butyrolactol A | Amphotericin B | Caspofungin | Fluconazole | Terbinafine |
| *Cryptococcus neoformans* H99 | 1~2 | 0.5 | 32 | 8 | 1 |
| *Cryptococcus neoformans* H99 (SDB) | 0.06~0.125 | 0.125 | 1~2 | 32 | 0.5 |
| *Cryptococcus neoformans* CDC15 | 2 | 0.5 | 32 | 16 | ND |
| *Cryptococcus neoformans* JEC20 | 2 | 0.25 | 16 | 8 | ND |
| *Cryptococcus gattii* R265 | 2 | 0.25 | 32 | 8 | ND |
| *Cryptococcus gattii* R272 | 1 | 0.5 | 32 | 16 | ND |
| *Cryptococcus gattii* WM276 | 1 | 0.25~0.5 | 32 | 8 | ND |
| *Candida auris* CBS10913 | 4 | 1 | 0.5 | 8 | ND |
| *Candida auris* CBS12372 | >64 | 1 | 0.5 | >64 | ND |
| *Candida auris* CBS12373 | >64 | 1 | 0.5 | >64 | ND |
| *Candida auris* CBS12766 | 8~16 | 4~8 | 64 | >64 | ND |
| *Candida auris* CBS12775 | 8 | 4~8 | 64 | >64 | ND |
| *Candida auris* CBS12776 | 16 | 4~8 | 64 | >64 | ND |
| *Candida albicans* ATCC 90028 | 1 | 0.5 | 0.25 | 8 | 2 |
| *Candida albicans* ATCC 90028 (+0.8 M sorbitol) | 1 | 0.5 | 16 | ND | ND |
| *Candida albicans* ATCC 200955$^{AmBR}$ | >64 | 2 | 0.5 | >64 | 64 |
| *Candida albicans* SN95 | 1 | 1 | 0.25 | 8 | ND |
| *Candida albicans* DPL15$^{CasR}$ | 2 | 1 | 2 | >64 | ND |
| *Candida albicans* DPL21$^{CasR}$ | 4 | 1~2 | 4 | 32 | ND |
| *Candida albicans pkc1/pkc1*$^{\Delta pkc1}$ | 1 | 0.5 | 0.125 | 2 | ND |
| *Candida parapsilosis ATCC90018* | 4 | 1 | 0.5 | 0.25 | ND |
| *Candida parapsilosis ATCC22019*$^{FluR}$ | 1 | 1 | 2 | 2 | ND |
| *Candida tropicalis* | 2 | 1 | 0.25 | 1 | ND |
| *Candida tropicalis ATCC200956*$^{AmBR}$ | 8 | 8 | 0.25 | >64 | ND |
| *Nakaseomyces glabratus*$^{FluR}$ | 4 | 1 | 0.5 | 4~8 | ND |
| *S. pombe ATCC38366* | 0.125 | 2 | 1 | 1 | ND |
| *S. pombe* | 0.125 | 0.5 | 1 | 1 | ND |
| *Saccharomyces cerevisiae DL1* | 1 | 0.25 | 0.125 | 2 | ND |
| *Saccharomyces cerevisiae BY4741* | 2~4 | 0.25 | 0.125 | 4 | ND |
| *Aspergillus fumigatus* Af293 | 4 | 0.5 | 64 | >64 | ND |
| *Trichophyton rubrum* ATCC28188 | 2 | 1 | 64 | 64 | ND |

Minimal inhibitory concentrations (MICs) of butyrolactol A were determined via broth microdilution against various fungal isolates, compared to established clinical agents. Superscript *AmBR*: Amphotericin B-resistant strains. Superscript *CasR*: Clinical echinocandin-resistant isolates. Superscript *Δpkc1*: *PKC1* gene deleted from SN95. Compound concentrations are presented in μg/mL.

**Supplementary Table 6**. **Potentiation activity of butyrolactol A to caspofungin against *Cryptococcus spp.* and multi-resistant *Candida auris*.**

| **Organism** | **MIC (μg/mL)** | | | | |
|---|---|---|---|---|---|
| | BLA | CAP | CAP (+ ½ MIC BLA) | Rescue Concentration | FICI(CAP+BLA) |
| *Cryptococcus neoformans* H99 | 2 | 32 | 1 | 1 | 0.375 |
| *Cryptococcus neoformans* JEC20 | 1~2 | 16 | 2 | 0.5 | 0.375 |
| *Cryptococcus gattii* R265 | 2 | 32 | 2 | 1 | 0.375 |
| *Cryptococcus gattii* WM276 | 1 | 32 | 1 | 0.5 | 0.375 |
| *Candida auris* CBS12766 | 8 | 32 | 0.125 | 2 | 0.25 |
| *Candida auris* CBS12776 | 16 | 64 | 0.125 | 2 | 0.125 |

The MIC of caspofungin (CAP) was determined both in the absence and presence of ½ MIC of butyrolactol A (BLA). MIC refers to the minimum inhibitory concentration, and compound concentrations are expressed in μg/mL. The Rescue Concentration is the concentration of BLA that reduces the MIC of CAP to the breakpoint (2 μg/mL).

**Supplementary Table 7**. Summary of cytotoxicity assessment of butyrolactol A.

| **Cell Line** | | **$IC_{50}$ (μg/mL)** | | |
|---|---|---|---|---|
| | Assay Used | Butyrolactol A | Amphotericin B | MG-132 |
| *HEK293* | ATP Viability | 285.9 | 30.59 | 0.3876 |
| *HEK293* | LDH Cytotoxicity Assay | 126.5 | 9.424 | 1.654 |
| *HepG2* | ATP Viability | 728.6 | 14.80 | 0.3662 |
| *HepG2* | LDH Cytotoxicity Assay | 44.60 | 1.749 | 0.9429 |
| *THP-1* | ATP Viability | 5.267 | 1.110 | 0.4711 |
| *THP-1* | LDH Cytotoxicity Assay | 4.405 | – | 0.5142 |

Note: the Butyrolactol A and Amphotericin B data represent the average of 6 values (triplicates from two separate days). No values are reported for amphotericin B as they could not be calculated from the curve.

**Supplementary Table 8**. **Cross-test between the BLA-resistant mutants and wild-type strain KN99a**

| Resistant Parent | Wild Type Parent | Resistant Parent Genotype | Total Progeny | Phenotype Segregation Resistant: Sensitive (%) | | Presence of parental *APT1* mutation in resistant progeny (%) | |
|---|---|---|---|---|---|---|---|
| | | | | *Observed* | *Expected* | *Observed* | *Expected* |
| $BLA^R$-RA3 mutant | WT KN99a | *apt1-2* (frameshift) | 26 | 14:12 (54%) | 13:13 (50%) | 14/14 (100%) | 14/14 (100%) |
| $BLA^R$-RA4 mutant | WT KN99a | *apt1-3* (G266V) | 19 | 9:10 (47%) | 9.5:9.5 (50%) | 8/9 (89%) | 9/9 (100%) |

**Supplementary Table 9**. Cryo-EM data collection, refinement and validation statistics.

| | Apt1–Cdc50 in E2P state bound with BLA (EMD-70618) (PDB 9OMV) | Apt1–Cdc50 in E1 state (EMD-47339) (PDB 9DZV) |
|---|---|---|
| **Data collection and processing** | | |
| Microscope | Titian Krios | Titian Krios |
| Camera | K3 | K3 |
| Magnification | 105,000 | 105,000 |
| Voltage (keV) | 300 | 300 |
| Electron dose ($e^-/Å^2$) | 60 | 60 |
| Exposure rate ($e^-$/px/s) | 41.1 | 41.1 |
| Number of frames | 50 | 50 |
| Slit width (eV) | 20 | 20 |
| Data acquisition software | SerialEM | SerialEM |
| Defocus range (-µm) | 1.3-1.7 | 1.3-1.7 |
| Pixel size (Å) | 0.828 | 0.828 |
| Symmetry imposed | C1 | C1 |
| Micrographs collected (no.) | 11,811 | 11,436 |
| Particle images used for 3D (no.) | 1,830,211 | 1,884,807 |
| Final particle images (no.) | 187,186 | 131,633 |
| Map resolution (Å) | 3.0 | 3.3 |
| FSC threshold | 0.143 | 0.143 |
| Map resolution range (Å) | 1.8-9.5 | 1.9-10.6 |
| cFSC Area Ratio (cFAR) | 0.52 | 0.33 |
| **Refinement** | | |
| Refinement package | Phenix | Phenix |
| Initial model used (PDB code) | AlphaFold *in silico* model | PDB 9OMV of this study |
| Map correlation coefficient (MapCCvolume/mask) | 0.80/0.79 | 0.79/0.79 |
| Model resolution (Å) | 3.1 | 3.3 |
| FSC threshold | 0.5 | 0.5 |
| Map sharpening *B* factor ($Å^2$) | -92 | -95.3 |
| Model composition | | |
| Non-hydrogen atoms | 11,594 | 11,502 |
| Protein residues | 1437 | 1431 |
| Ligands | 14 | 12 |
| *B* factors ($Å^2$) | | |
| Protein | 66.51 | 81.77 |
| Ligand | 56.60 | 75.89 |
| R.m.s. deviations | | |
| Bond lengths (Å) | 0.003 | 0.003 |

| | | |
|---|---|---|
| Bond angles (°) | 0.522 | 0.524 |
| **Validation** | | |
| MolProbity score | 2.34 | 2.63 |
| Clashscore | 8.85 | 12.36 |
| Poor rotamers (%) | 3.74 | 4.33 |
| CaBLAM outliers (%) | 3.69 | 7.04 |
| EMRinger score | 3.07 | 2.68 |
| Average Q-score | 0.44 | 0.40 |
| Ramachandran plot | | |
| Favored (%) | 93.46 | 90.27 |
| Allowed (%) | 6.54 | 9.73 |
| Disallowed (%) | 0.00 | 0.00 |

**Supplementary Table 10**. Oligonucleotides used in the study

| REAGENT or RESOURCE | SOURCE | IDENTIFIER |
|---|---|---|
| oligonucleotides | | |
| VIntron1-F: GTGTTGAGGTGGGAGAGAC | This paper | N/A |
| VIntron1-R: GGGAAGATGTCAAGGTTGG | This paper | N/A |
| Donor DNA3-F: GGGCCAGGCTTAAGATTC | This paper | N/A |
| Donor DNA3-R: GGGTTCTAGGATTGAAATGG | This paper | N/A |
| CXF-APT13end-VF1: CTGAGCAAGGATCTCTTCTTC | This paper | N/A |
| CXF-exconst-VR1: GAAGGTATCATGAACGTCCG | This paper | N/A |
| Vector FP: ATTCCAAGAATGTGAGCTCTTAATTAACAATTCTTCGC | This paper | N/A |
| Vector RP: AATGGTTTGTAATAAGATCCGCTCTAACCGAAAAG | This paper | N/A |
| GAL FP: GAAGATAGCCATGGGGTTTTTCTCCTTGACG | This paper | N/A |
| GAL RP: TTTGGAGGCACCCTTATCGTCGTCATCCTTGTAATCC | This paper | N/A |
| N1109A FP: TCTGGGCTGTTTTCTGGACCTTG | This paper | N/A |
| N1109A RP: AGAGCAAGTAAACGTAAGCGAAGACGTAG | This paper | N/A |
| P127A FP: GGTTATCTTGGCTTTGATTATTGTTTTGGC | This paper | N/A |
| P127A RP: AAACCTGGAGAAATAGTAGAGAACTTTGG | This paper | N/A |

**Supplementary Note 1. NMR data for Butyrolactol A and B in dmso-d6.**

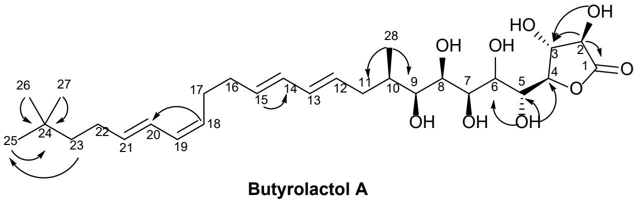


**Butyrolactol A**

Selected HMBC interactions

| | Butyrolactol A | |
|---|---|---|
| Carbon # | $\delta_H$ (m, J in Hz) | $\delta_C$ |
| 1 | - | 175.21 |
| 2 | 4.26 (dd, *J* = 9.0, 6.5 Hz, 1H) | 74.03 |
| 2-OH | 6.04 (d, 1H, 6.4) | - |
| 3 | 4.16 (td, J = 8.7, 6.1 Hz, 1H) | 72.38 |
| 3-OH | 5.70 (d, *J* = 6.2 Hz, 1H) | - |
| 4 | 4.32 (dd, *J* = 18.0, 8.2 Hz, 1H) | 79.63 |
| 5 | 3.63 (t, *J* = 8.7 Hz, 1H) | 66.38 |
| 5-OH | 4.85 (d, *J* = 7.7 Hz, 1H) | - |
| 6* | 3.72 (dt, *J* = 10.6, 8.5 Hz, 1H) | 68.47 |
| 6-OH | 4.32 (m,1H) | - |
| 7* | 3.72 (dt, *J* = 10.6, 8.5 Hz, 2H) | 68.32 |
| 7-OH | 4.02 (d, *J* = 8.2 Hz, 1H) | - |
| 8 | 3.49 (t, *J* = 8.9 Hz, 1H) | 69.14 |
| 8-OH | 3.97 (d, *J* = 8.2 Hz, 1H) | |
| 9 | 3.36 (d, *J* = 8.7 Hz, 1H) | 72.52 |
| 9-OH | 3.93 (d, *J* = 8.4 Hz, 1H) | - |
| 10 | 1.69 (dtt, *J* = 9.4, 6.7, 2.9 Hz, 1H) | 35.65 |
| 11a,b | 1.86 (dt, *J* = 13.9, 8.5 Hz, 1H)<br>2.44 (ddd, *J* = 14.6, 6.6, 3.0 Hz, 1H) | 35.74 |
| 12 | 5.56 (ddt, *J* = 21.7, 14.4, 7.1 Hz, 1H) | 131.38 |
| 13 | 5.98 (m, 1H) | 131.43 |
| 14 | 6.04 (m, 1H) | 130.84 |
| 15 | 5.56 (ddt, *J* = 21.7, 14.4, 7.1 Hz, 1H) | 131.45 |
| 16 | 2.11 (m, 2H) | 32.09 |
| 17 | 2.21 (q, *J* = 7.4 Hz, 2H) | 27.03 |
| 18 | 5.27 (dt, *J* = 10.9, 7.5 Hz, 1H) | 128.65 |
| 19 | 5.93 (t, 1H, 11.1) | 128.98 |
| 20 | 6.30 (dd, *J* = 15.3, 10.7 Hz, 1H) | 125.19 |
| 21 | 5.66 (dt, *J* = 14.5, 7.0 Hz, 1H) | 135.35 |

| | | |
|---|---|---|
| 22 | 2.01 (m, 2H) | 27.71 |
| 23 | 1.26 (m, 2H) | 43.0 |
| 24 | - | 30.16 |
| 25 | 0.87 (s, 9H) | 29.08 |
| 26 | | |
| 27 | | |
| 28 | 0.77 (d, $J = 6.7$ Hz, 3H) | 15.51 |

*overlapping signals

$^{1}$H- NMR spectrum of butyrolactol A.

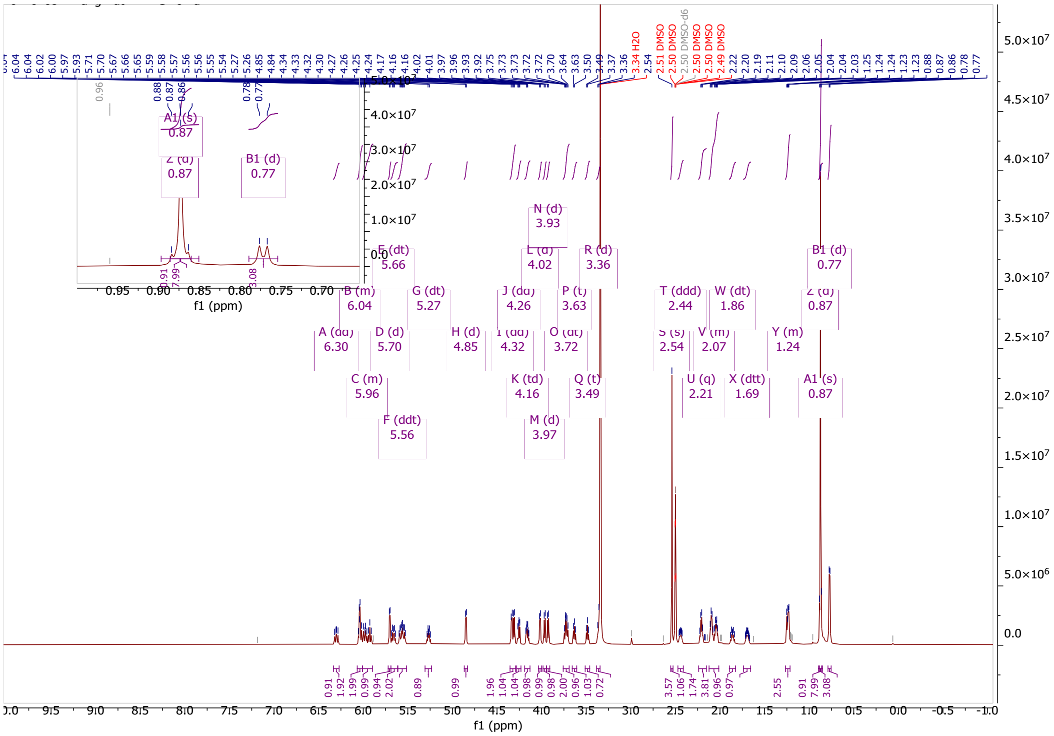

$^{13}$C-deptq NMR spectrum of butyrolactol A.

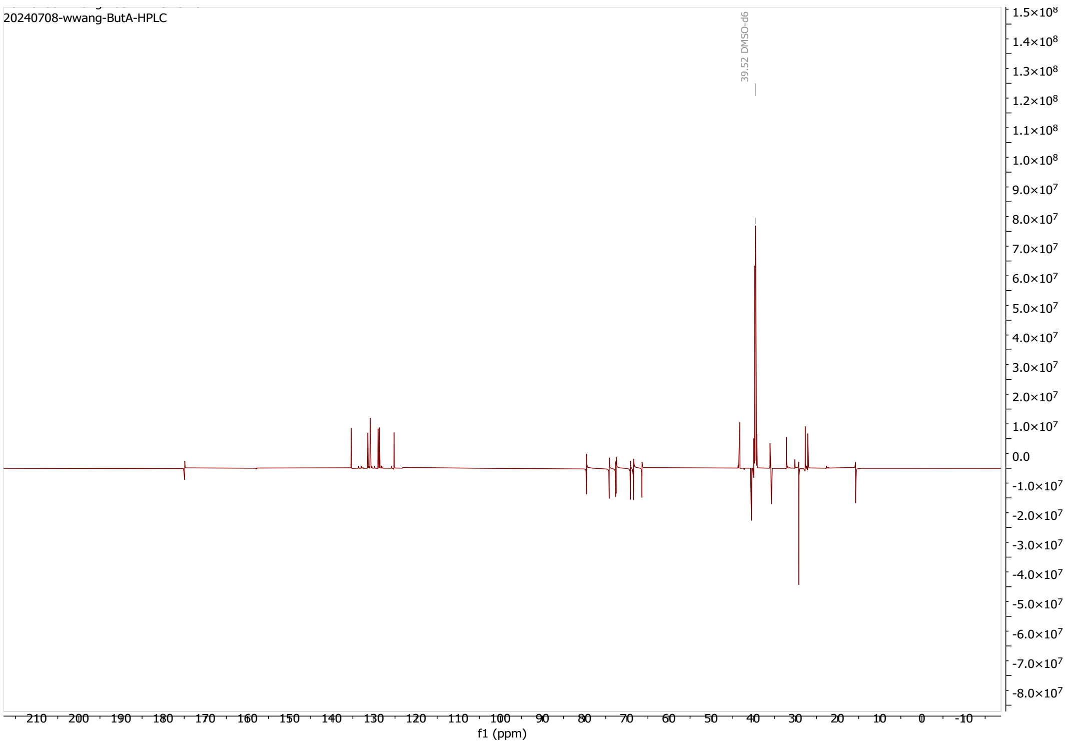


$^{1}$H-$^{1}$H- COSY NMR spectrum of butyrolactol A.

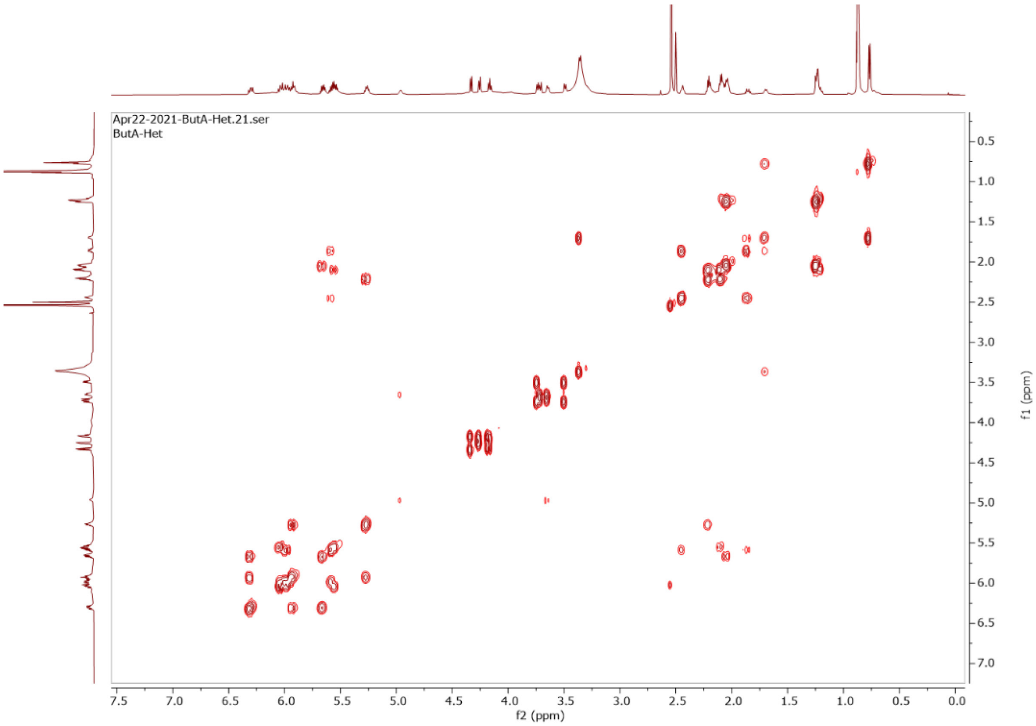

$^{1}$H-$^{13}$C-NMR- HSQC spectrum of butyrolactol A.

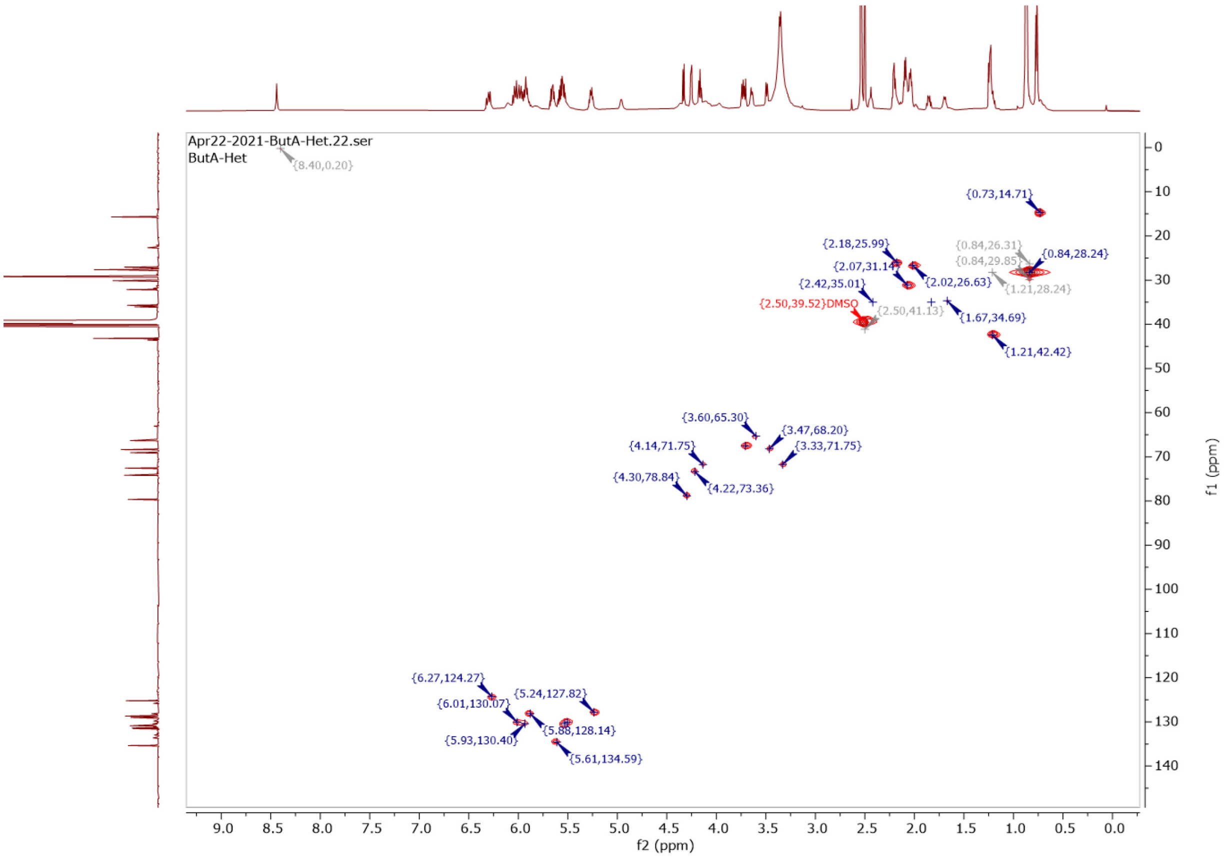


**Supplementary Note 2. Assignment of the relative and absolute configurations in butyrolactol A (BLA)**

To determine the stereochemistry of butyrolactol A (BLA), we performed detailed NMR-based structural analysis combining vicinal $^{3}J$ $^{1}$H–$^{1}$H and long-range $^{2,3}J$ $^{1}$H–$^{13}$C coupling constants with ROESY correlations.[2] For the γ-lactone–polyol head group, large coupling constants between H-2/H-3 (8.6 Hz) and H-3/H-4 (8.5 Hz) indicated anti-periplanar orientations and supported a 3R,4S configuration (Supplementary Note 1).

In the extended polyol tail, the absence of a detectable coupling between H-4 and H-5 (0 Hz) suggested an orthogonal dihedral angle, and ROESY correlations between H-3/H-5, H-3/5-OH, and H-5/3-OH supported a 5R configuration. Strong coupling constants between H-5/H-6, H-7/H-8, and H-9/H-10 (8.9, 9.3, and 9.0 Hz, respectively), along with diagnostic ROESY cross-peaks, indicated erythro relationships and anti orientations across these segments. C6 was assigned as 6R.

Smaller couplings (<3 Hz) between H-6/H-7 and H-8/H-9 indicated gauche conformations, while $^{2}J$ $^{1}$H–$^{13}$C couplings (1.7 Hz and 2.8 Hz) from HSQC-HECADE supported a threo configuration at

C8–C9. These data collectively supported the relative configuration from C7 to C10 as 7S*, 8S*, 9R*, and 10R*.

In the conjugated diene regions, olefinic coupling constants between H-12/H-13 (14.4 Hz), H-14/H-15 (15.0 Hz), H-18/H-19 (11.0 Hz), and H-20/H-21 (15.0 Hz) were consistent with E geometry at C12=C13, C14=C15, and C20=C21, and Z geometry at C18=C19.

A summary of stereochemical assignments and supporting NMR spectra is provided in Supplementary Notes 1 and 2.

HETLOC NMR spectrum of butyrolactol A.

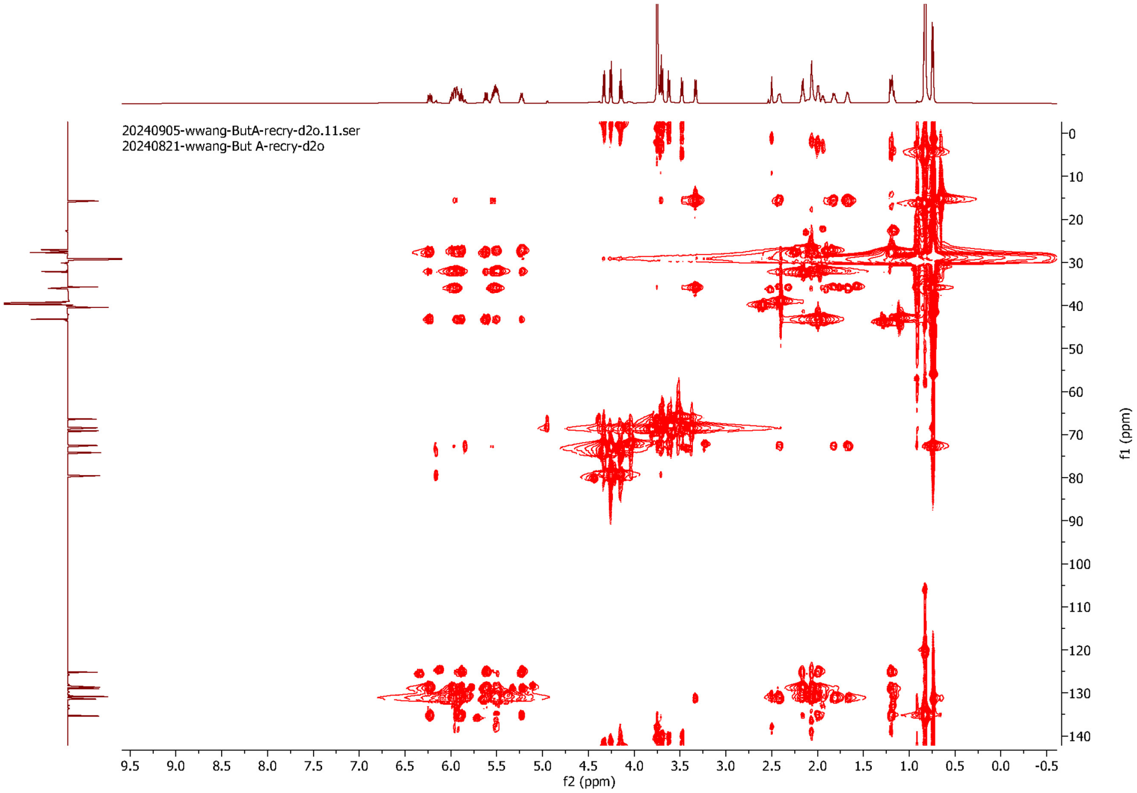

ROESY NMR spectrum of butyrolactol A.

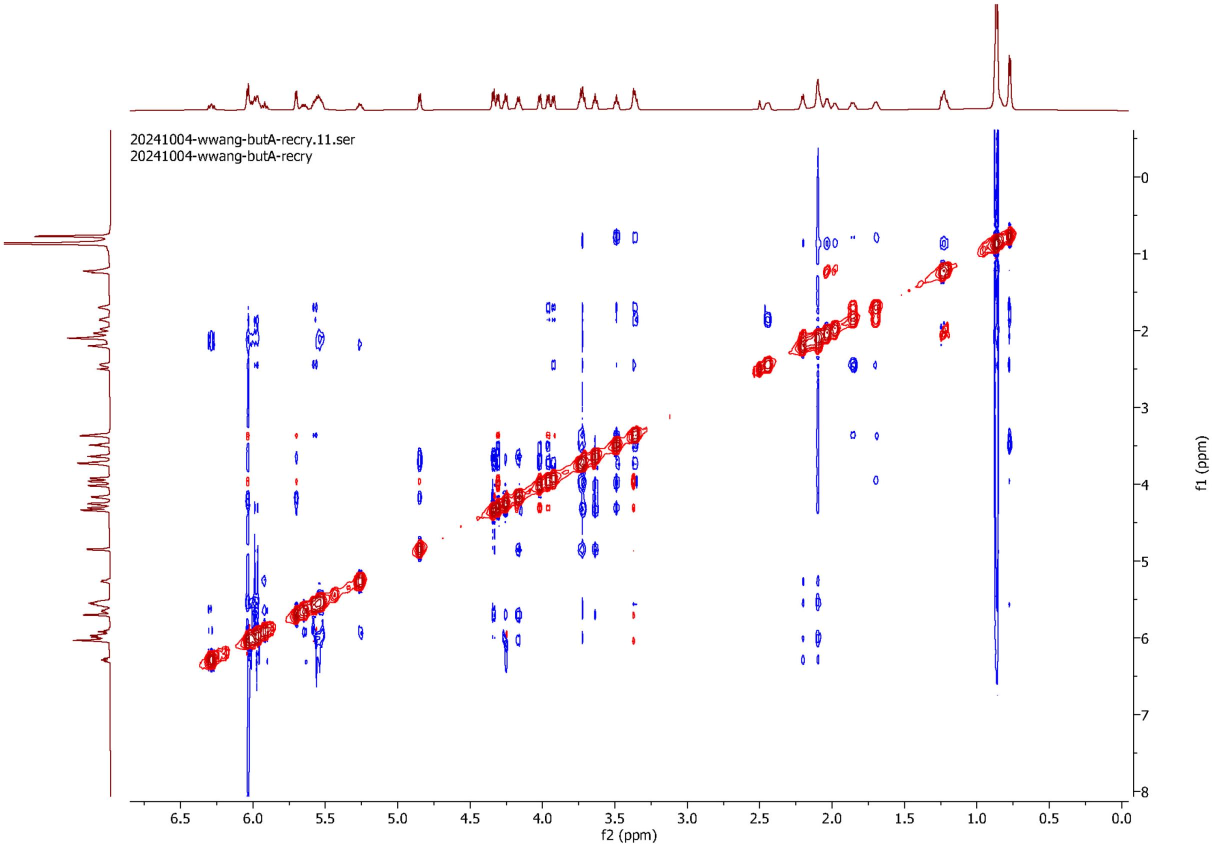
